# Rapid Changes in Risk Attitudes Originate from Bayesian Inference on Parietal Magnitude Representations

**DOI:** 10.1101/2024.08.23.609296

**Authors:** Gilles de Hollander, Marcus Grueschow, Franciszek Hennel, Christian C. Ruff

**Author notes:** **Corresponding authors:** Gilles de Hollander; Christian C. Ruff. **Competing Interests** The authors declare no competing interests.

## Abstract

Risk attitudes – the willingness to accept greater uncertainty to achieve larger potential rewards – determine many aspects of our lives and are often interpreted as an individual trait that reflects a general ‘taste’ for risk. However, this perspective cannot explain why the choices that people make concerning risky outcomes can change considerably across contexts and even across repetitions of the identical decisions. Here we provide modelling and neural evidence that such rapid changes in risk attitudes emerge from Bayesian inference on noisy magnitude representations in the parietal cortex. Our participants underwent fMRI while choosing between safe and risky options that were either held in working memory or present on the screen. Risky options that were held in working memory were less likely to be chosen (risk aversion) when they had relatively large payoffs but more likely to be chosen (risk-seeking) when they had relatively small payoffs. These counterintuitive effects can be explained mechanistically by a computational model of the Bayesian inference underlying payoff perception: Options kept in working memory are noisier and therefore more prone to central tendency biases, leading small (or large) payoffs to be overestimated (or underestimated) more. Congruent with the behavioural modelling, fMRI population-receptive field modelling showed that on trials where intraparietal payoff representations were noisier, choices were also less consistent and less risk-neutral, in line with participants resorting more to their prior beliefs about potential payoffs. Our results highlight that individual risk attitudes and their puzzling changes across contexts and choice repetitions can be rooted in perceptual inference on noisy parietal magnitude representations, with profound implications for economic, psychological, and neuroscience theories of risky behaviour.

## Introduction

People differ widely in their attitudes towards risk. In lab experiments, some people will forgo a lottery with a larger expected value for a substantially lower but certain payout, whereas others will choose the lottery^1–9^. Relatedly, in everyday life, some people are more willing than others to become self-employed, engage in new relationships, or to invest money in the stock market^10^. Neoclassical economic theories describe such individual differences in behavior using the concept of individual risk preferences: An individual would be *risk-averse* if they prefer to obtain for sure the expected value of a risky prospect (its gain multiplied by its probability) to realizing the risky prospect itself, *risk-seeking* if they prefer the risky prospect itself to obtaining its expected value for sure, and *risk-neutral* if they are indifferent between these options. These preferences are believed to be (relatively) stable and generalizable across choice domains^3,4,11^. Preferences are by definition latent and cannot be measured directly; however, they can be inferred from overt behavior (choices “reveal” preferences, in the terminology of neoclassical economics). Thus, measuring overt risk attitudes in laboratory choices is the standard way to infer underlying risk preferences, which is seen as key for any statements, recommendations, or interventions designed to help people take decisions under uncertainty in a way that increases their chances to meet personal goals.

Empirical data reveal, however, that individual risk attitudes are highly unstable^12,13^. Objectively identical choice sets (i.e., accept/reject specific gambles) can lead to different choices depending on the options that were offered before^14^ or in which order choice options are presented^15^. Furthermore, even when the exact same choice problem is presented repeatedly within the same context, decision-makers choose differently across repetitions^7,16,17^. This seems at odds with the idxea that these choices mainly reveal stable preferences.

Adaptations of the preferential framework have tried to account for the stochasticity of risky choice by adding a random noise term to the utility function (Random Utility Theory, RUT)^18^ or by assuming that decision-makers may have a preference for responding randomly^19^. Although these theories alleviate the problem of choice inconsistency, they do not address the question of where this randomness comes from, or why risk attitudes are so dependent on context (e..g, order of presentation). Both theories frame the noise in observed behaviour as random and therefore external to the decision-making processes dealing with the choice-relevant information (e.g., payoffs). This stands in stark contrast to numerous findings in perceptual neuroscience that the degree of randomness in observed choices is usually tightly linked to the amount of bias in decisions, suggesting a common origin^20–26^.

In contrast to RUT, more recent economic choice models attribute risk attitudes directly to the noisiness of how decision-makers perceive the choice problems they are faced with^27–31^. These perceptual models of risky choice draw on theories from perceptual neuroscience, framing decisions as fully rational solutions to optimization problems computed on inherently noisy representations of choice-relevant information. To make optimal choices, our brain makes the best of these noisy signals by applying principles of Bayesian inference^24,32^ and/or efficient coding^26,33^, which can lead to biases in percepts of the lottery payouts and probabilities. Note that, on average, such biases are beneficial because they reduce the average perceptual error, by employing prior information about the likely causes of sensory signals in case these are noisy and therefore unreliable. For example, there is a well-known positive correlation between the levels of noise in cognitive representations and central tendency (CT) bias^34^: With increased noise, participants tend to regress their percepts of stimulus values towards the mean value of earlier observed stimuli. The Bayesian interpretation of this is that, when noise increases, participants also increasingly revert to their prior beliefs about possible stimulus values – just as normative statistical theories prescribe^16,24^.

In the context of risky choices, decision-makers with noisier perception of a choice problem therefore tend to estimate the larger (risky) payoffs downward towards a prior mean, but the smaller (safe) payoffs upwards. As a consequence, decision-makers may underestimate the size of the larger risky payoffs, thereby exhibiting more risk-averse attitudes. This suggests that two people might in fact be similarly risk-neutral, but may show substantially different risk attitudes in their choices simply because they differ in how precisely and thus unbiased they can perceive a risky choice problem.

We have recently provided initial neuroscientific evidence for such a perceptual account of risk attitudes, by showing that there is a link between the noisiness of an individual’s neurocognitive magnitude representations in parietal cortex during perceptual judgments and the amount of risk aversion they exhibit in separate economic choices^35^. This link between neural measurements and latent cognitive variables paves the way for asking important questions about the neurocognitive origins of changes in risk attitudes. First, what determines whether individuals show categorically distinct risk attitudes such as risk-seeking, risk-neutral, and risk-averse behaviour^5^? These categories currently still elude perceptual models of risky choice^27,30^, which have until now focused on risk aversion. Second, risk attitudes appear highly sensitive to contexts: Choices can differ substantially depending on how decision problems are framed^4^, in which order options are presented^15^, or which choice options were seen in earlier choice problems^14^. Can we mechanistically account for the influence of context on economic choice, by studying how it alters the perception of the choice-relevant information? Finally, as alluded to above, economic choice behaviour is profoundly variable even without any changes in context^7,16,17^. Where do these apparent fluctuations in risk attitudes come from? Can they originate from endogenous fluctuations of neural noise, in analogy to how such fluctuations affect perceptual decisions about low-level visual features^36^?

To address these questions, we introduce a general computational model of choice, the *Perception and Memory-based Choice Model (PMCM)*. The model builds upon existing perceptual models of risky choice^27,35^ that frame risky choice as a Bayesian inference process on noisy percepts of potential payoffs. Critically, in the PMCM, the noisiness of the different percepts relevant to the choice is no longer fixed but depends on the particular context of a specific trial. Specifically, in the experiment presented here, the noisiness of the representation of a choice option may depend on both whether a choice option is perceived directly or held in working memory, as well as on the acuity of neural processing on a particular trial. Thus, the PMCM makes it possible to characterize the psychological and neural mechanisms by which both exogenous environmental contexts and endogenous (neural) factors can shape economic choice on a moment-to-moment basis. This enables us to provide a decisive test of perceptual theories of risky-choice mechanisms using biological data.

In the following, we will first describe the risky choice paradigm we developed to modulate the acuity with which choice options are represented in the brain, by manipulating the order in which they are presented. This experimental paradigm also allows us to extract an fMRI signal that corresponds specifically to the neural acuity of the perception of the payoff of the first choice option, independent from any choice-related neural activity that comes later in the trial^35^. To be able to do so, we used the numerical magnitudes of dot clouds to present the payoff magnitudes to the participants, as we have in earlier work^35^. Such stimuli are somewhat different from the Arab numerals often employed to present potential payoffs; however, seminal work in numerical cognition has established that the neural processing of the numerosity of such stimuli is similar to that of more general (symbolic) numerical information^37,38^, including Arab numerals^35^. Numerosity processing for both types of stimuli engages a more general parietal ‘magnitude system’^39^, and the acuity of numerosity processing for both types of stimuli is correlated across individuals^35^.

We will then introduce in detail the PMCM and show how it can explain why some people are generally risk-seeking while others are risk-averse, as well as why people appear risk-averse for some sets of choices but risk-seeking for others. Finally, we will show that random fluctuations of sensory neural noise - the acuity of neural magnitude representations in the right intraparietal cortex as measured using inverted encoding models^35,40,41^ - relate mechanistically to choice consistency and risk attitudes in tandem, in line with a Bayesian perceptual account of risky choice. Our results provide evidence that risk attitudes and their variability across trials and contexts originate (at least partially) from aspects of perceptual neural processing in parietal cortex. These links, and thus the neurocognitive mechanisms underlying risk attitudes, can be captured by our Bayesian perceptual model of risky choice, which is not possible with classical models based on expected utility. Our approach and findings have profound implications for economic, psychological, and neuroscience theories of risk-taking, as well as the experimental paradigms used to study and evaluate risk attitudes in theoretical and applied contexts.

## Results

### Experimental paradigm

Thirty participants (14 female, aged 20-34) performed the task twice: Once in a 3 Tesla (T) scanner and once in a 7T scanner. We chose to acquire data on two field strengths to allow for both eye-tracking (3T; data not presented here) as well as ultra-high resolution imaging (7T). To study how risk attitudes relate to the acuity of perception and the underlying neural representations, we developed a new experimental paradigm based on earlier work^35^. In our paradigm, participants had to choose between a sure payoff and a gamble with a 55% probability of winning a larger payoff (Fig. 1A). Crucially, the payoffs of the safe and risky options were presented sequentially; in some trials, the risky options were presented first, in other trials, the safe option was presented first. This design aimed to introduce working memory noise for the first-presented option that had to be memorized, thereby presumably enhancing CT biases for this option.

**Figure 1:**
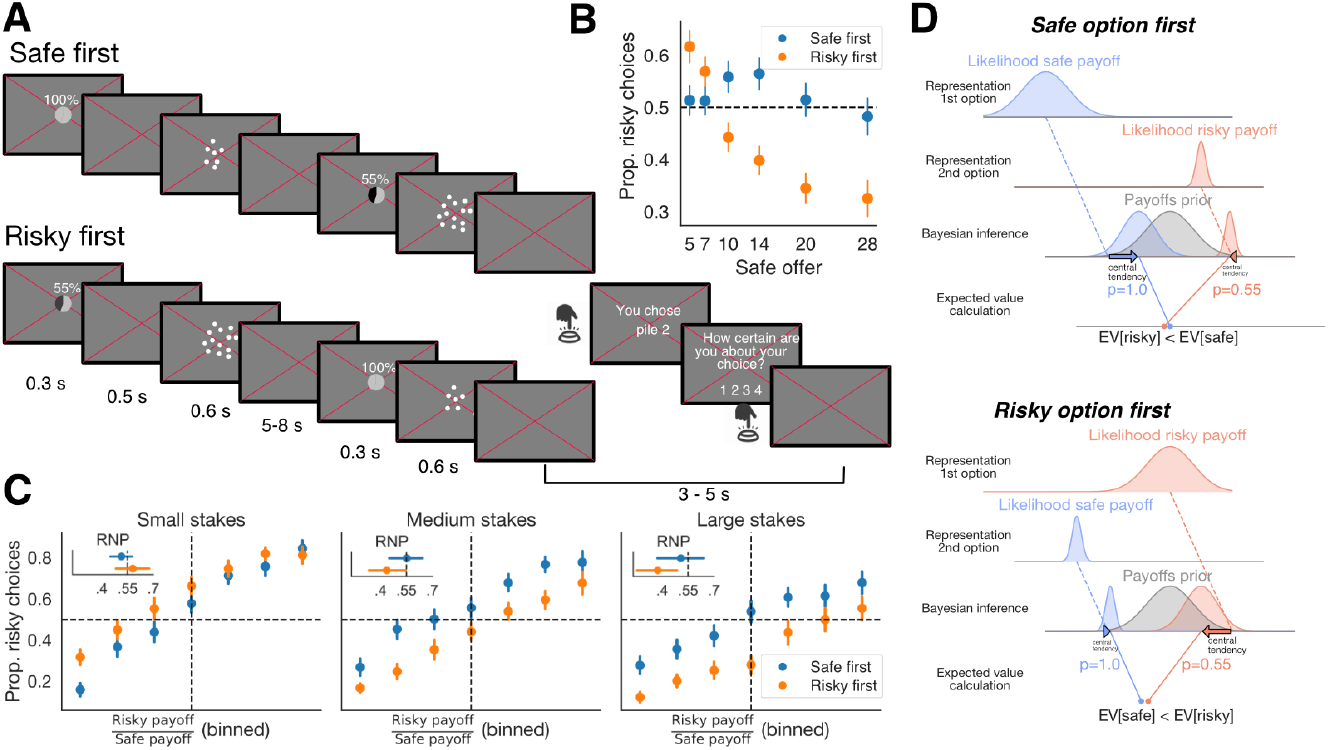
**A)** Participants were incentivized to choose between two prospects: A risky option with 55% probability of receiving the payoff and a safe option with 100% probability of receiving the payoff. Prospects were presented sequentially as clouds of 1-CHF coins, and payoff probabilities were presented as both piecharts and symbolic percentages just before the payoff information. Crucially, either the risky or the safe option was presented first and thus had to be held in working memory (all choice sets occurred twice for every order for every session). **B)** Average proportion of risky choices (y-axis) as a function of the magnitude of the safe option (x-axis) and presentation order (hue). Participants were either more or less likely to choose the safe option when the risky option was presented first, depending on whether the stake size (i.e., indexed by the magnitude of the safe option, around which the magnitude of the risky option was varied) was larger or smaller. Error bars correspond to the standard error of the mean over participants. **C)** Psychophysical curves for different stake sizes (defined as the magnitude of the safe option), i.e., the probability of a risky choice as a function of the ratio between risky and safe payoffs (binned per participant based on their calibration data, i.e., vincentization). Insets show the estimated risk-neutral probability, a measure of average risk attitude (RNP<55% = risk-averse, RNP>55% = risk-seeking). For relatively small-stakes gambles, participants become (more) risk-seeking when the risky option is presented first. Conversely, for relatively large-stake gambles, they became more risk-averse. This is in line with larger central-tendency effects for remembered versus currently presented payoffs. **D)** Illustration of the intuition behind the Perception and Memory-based Choice (PMC) model: When an option is presented first, its representation will be noisier because the option needs to be held in working memory and the participant has no direct access to the corresponding stimulus during the decision (unlike the second option). Thus, the very same risky option has a noisier likelihood when presented first (left panel) as compared to when it is presented second (right panel). The first option will therefore be more influenced by the prior than the second option and will have a larger CT bias. Presentation order can therefore impact on risk attitude in different ways, depending on stake size. Note that this illustration uses plausible but stereotyped parameter values for visualization purposes.

We kept the payout probability of the risky option fixed at 55%, in line with previous studies^27,35^, to streamline the computational modeling of behavioral and neural data. While we recognize the general significance of probability distortion in decision-making^42^, our study focussed on the perceptual impacts of payoff evaluations. This focus required a design that emphasized statistical power for these specific effects, and thus fixed payoff probabilities.

The safe option consistently offered one of six possible payoffs, with equal probability but in random order: 5, 7, 10, 14, 20, or 28 CHF. The risky option was determined based on one of eight possible ratios relative to the safe option. These ratios were established through a calibration task conducted outside the scanner, ensuring that the choices presented to participants were non-trivial, as they were near the participants’ individual indifference points (see Methods for details). Crucially, the payoffs of the safe and risky options were presented sequentially, with in some trials the risky options being presented first, and in other trials the safe option being presented first. This design aimed to introduce working memory noise for the first-presented option that had to be memorized, thereby presumably enhancing CT biases for this option.

In our fMRI experiment, the payoffs were presented as dot clouds (rather than Arabic numerals) because previous work has indicated that such stimulus magnitudes can be reliably decoded from rIPS^35,43^. However, note that we have also performed a very similar experiment using Arabic numerals outside of the scanner (see Supplementary text S3 for more details).

### Different noise contexts shift risk attitudes as predicted by Bayesian inference on noisy payoff representations

We first examined how exogenous factors that modulate neurocognitive noise – in our case, the presentation order of the options – impacted choice behaviour. Psychophysical work on the noisiness of working memory^44–46^ suggests that the neurocognitive representation of the initially presented payoffs, retained in working memory, are subject to more noise compared to the options presented subsequently. Thus, taking a perceptual Bayesian perspective^16,24,27,35^, we predicted that the options that were presented first should be more sensitive to the central tendency (CT) bias, resulting in overestimation of relatively small payoffs and underestimation of relatively large payoffs. As a consequence, when a relatively small risky payoff was presented first, participants should be more likely to choose it compared to when it was presented second. Conversely, when a relatively large risky option was presented first, it should be less likely to be chosen. For safe options that were presented first, we expected an analogous pattern - in other words, risk attitudes should be categorically changed by presentation order and overall stake size.

To quantify risk attitudes, we used a psychophysical (probit) model which estimates the indifference point—the ratio between risky and safe payoffs at which individuals show no preference^27,35,47^. Critically, this model disentangles choice consistency from preference, avoiding the confounds inherent in raw choice proportions^47^. The indifference ratio can be converted to a *risk-neutral probability (RNP)*^*27*^, which corresponds to the probability for which a risk-neutral decision-maker would show the same indifference point. Thus, for our paradigm, a RNP of 55% corresponds to risk-neutral behavior, a RNP lower (or higher) than 55% to a risk-averse (or risk-seeking) attitude. In line with our predictions, our psychophysical modeling showed a strong interaction effect between order and overall stake size on RNP,. Specifically, formal model comparisons revealed that the full interaction model robustly outperformed simpler models with a model weight 95% using model stacking. Moreover, the expected log pointwise predictive density, ELPD^48^ (−5864.7), was at least 22.9 standard errors larger than for the other models; the next best model included only a main effect of stake size (ELPD −6121.4). In line with a CT effect for the first option, when stake sizes were small and the risky option came first, participants were, on average, risk-seeking (average RNP of 58.0% for lowest stakes, 95%CI 48.2-67.3%; see insets Fig. 1C) but became increasingly risk-averse as the average stake size increases (average RNP of 37.3% for highest stakes, CI 25.3-48.6%; there is a significant difference in RNP between the two presentation orders for all stake sizes, p_bayesian_<0.05)..

### The Perception and Memory-based Choice Model (PMCM)

To account for more complex risky choice environments, like the one in our experiment, we developed a novel perceptual model of (risky) choice. The model builds on classic psychophysical models of number perception^49,50^ as well as on economic models incorporating these insights to describe risky choice as a perceptual inference process, showing that risk aversion can arise as a result of central tendency effects in perception of the potential payoffs^27,35^. The crucial new feature compared to earlier models is that the acuity with which the payoffs of different choice options are represented can dynamically change as a function of exogenous (i.e., choice architecture) or endogenous (i.e., neural/physiological state) context. For example, in our particular experiment, the perception of the first- and second-presented payoffs can be differentially precise, as the noisiness of the neurocognitive payoff representation kept in working memory can be expected to be higher than the representation of the currently perceived payoff^46^. Furthermore, we can also use physiological indices of brain state (e.g., an fMRI activity decoding algorithm) to study how endogenous noise impacts choice behavior. A second novelty of our model is that it allows for different prior expectations about specific choice options based on secondary cues. For example, in most risky choice experiments, the payoffs of risky options are, on average, substantially higher than those of safe options and, given noisy perception, participants could use this information to their benefit. Note that both these features of our model and experiment resemble real-life choice situations, where options are rarely perceived exactly simultaneously and are almost always considered sequentially. By explicitly accounting for this in our design and model, we could test whether and how participants’ decisions are affected by this feature of a typical choice situation.

The full PMC model we employed in this study has six parameters: (1) the noisiness with which the first option is perceived (*ν*_1_), (2) the noisiness with which the second option is perceived (*ν*_2_), (3) the mean of the prior on risky payoffs (µ_*x*_), (4) the mean of the prior for the safe payoffs (µ_*c*_), (5) the standard deviation of the prior on risky payoffs (σ_*x*_), and (6) the standard deviation of the prior on safe payoffs (σ_*c*_). Note that these six parameters can be estimated in our experimental paradigm because the noise of the evidence is experimentally manipulated by order, orthogonal to the riskiness of the option. We confirmed the recoverability of the parameters of our model using multiple strategies. When the model was fitted separately to the same participants’ data for the two scanner sessions, the six parameters were all correlated across the two sessions, with correlation coefficients r(29) between 0.41 and 0.76 (all p<0.05). A parameter recovery study using the estimated parameters as generating parameters showed substantial correlations between generating and recovered parameters, ranging from 0.74 to 0.93. See Supplementary Text 1 for more details.

Posterior predictive plots confirmed that our model does an excellent job at predicting the observed empirical patterns at the group level, including the interaction between magnitudes and order of presentation (Fig. 2A). Moreover, our model also explains highly divergent behaviour across individual participants: Some participants are not influenced by order and/or overall stake sizes, and some participants are risk-seeking rather than risk-averse. Supplementary Fig. 1 gives some examples of typical behavioural patterns of representative participants and their posterior predictive plots.

**Figure 2:**
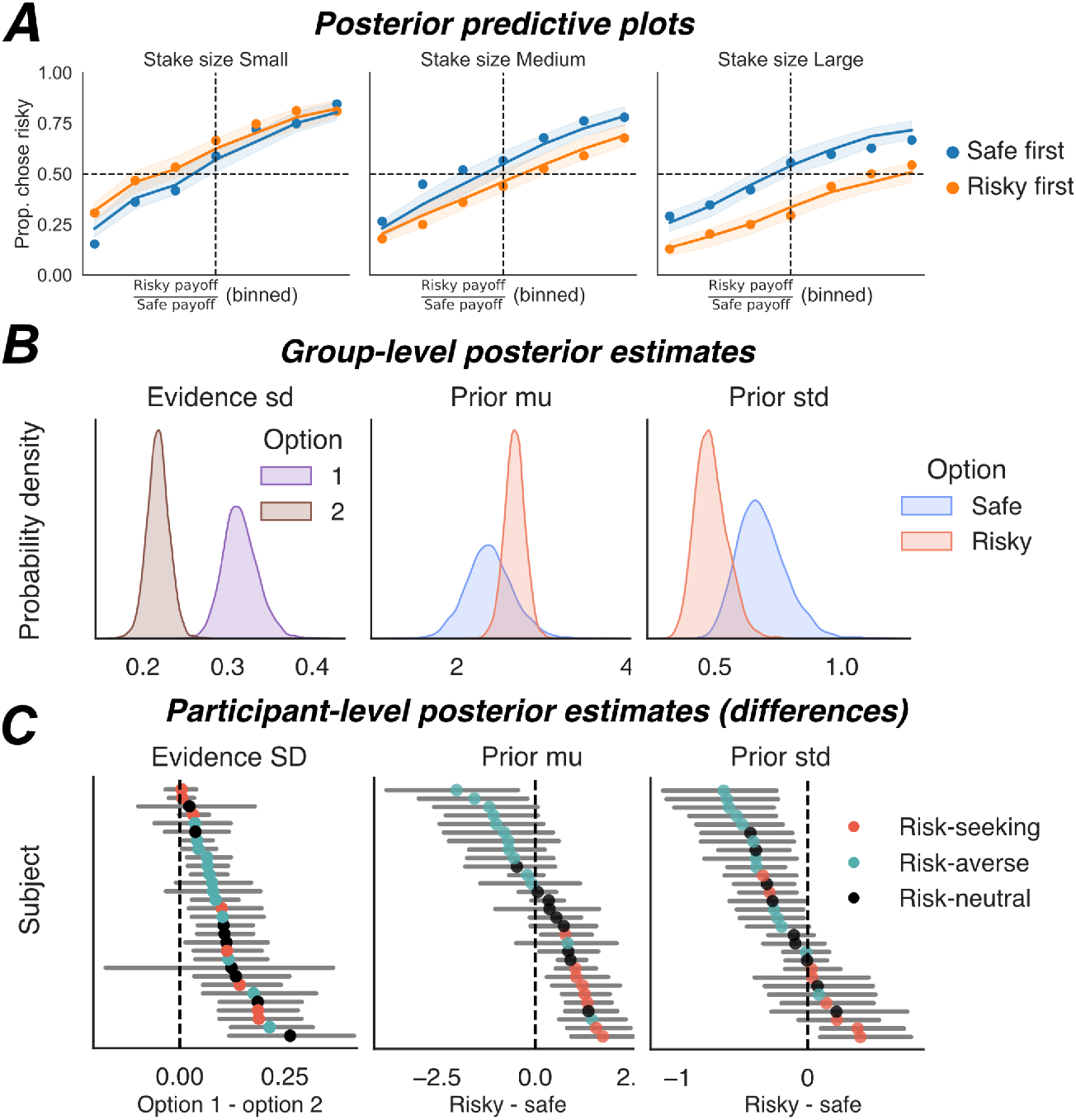
**A)** The PMC model’s posterior predictions (95% credible interval, shaded) closely match empirical data (markers), demonstrating an excellent fit. **B)** The estimated group parameter posteriors of the extended model suggest that the payoff of the first option is encoded with substantially more noise than the second option (left panel). Furthermore, the prior for risky options is estimated to have a higher mean payoff (middle panel; although this difference is significant only for some participants), as well as a more narrow range (right panel). Thus, risky payoffs are more prone to CT effects than safe payoffs. **C)** Participant-wise estimates of the differences between key parameters in the model, i.e., difference in noise between options presented first/second (left panel) and priors for the the safe and risky prior (middle/right pane). Dots are mean posterior estimates and lines are 95% credible intervals. Almost all participants have significantly ( p_Bayesian_< 0.05) noisier evidence about the first-presented payoff (leftmost panel), but only a subgroup employs different priors for risky and safe options (middle/right panel). This mean difference in risky and safe priors relates to the risk attitude of the participants (i.e., Risk-seeking/risk-averse or risk-neutral). For this figure, data from the 3T and 7T sessions were combined; see Supplementary Fig. 2 for separate analyses of the two sessions.

The first goal of our study was to explain how seemingly categorically different risk attitudes (e.g., risk aversion and risk seeking) may emerge from noisy perceptual inference processes. Our data reveal that there is indeed considerable variability across both participants and choice contexts when it comes to whether behaviour is ‘risk-seeking’, ‘risk-neutral’, or ‘risk-averse’. Indeed, we found that under some circumstances, our experimental population was risk-seeking on average, namely when the risky option was presented first and the safe option was relatively small (5 or 7; Fig. 1C). Our model-based inference that participants are particularly noisy and/or optimistic about potential prospects can now characterize the latent decision mechanisms and how they mechanistically lead to seemingly different behaviour in specific contexts. This appears more useful than a purely descriptive categorization based on average risk attitude.

The second main goal of our study was to explain why risk attitudes can shift across different contexts. Our model can naturally explain such context-specific risk attitudes: The risky payoff is particularly overestimated when it is relatively small and noisy. Moreover, it is well-known that a non-negligible fraction of experimental populations shows risk-seeking behaviour on average^5,51^. Current perceptual models of risky choice cannot account for this without assuming additional distortions in probabilities^27,30^. In contrast, our PMC model can account for these effects within one overarching framework: Risk-seeking participants have particularly optimistic prior beliefs about risky payoffs and are particularly risk-seeking when their percepts of the objective risky payoff are noisier (see Fig. 2C; left and middle panel). One question about our model might be whether it is unnecessarily complex, as it contains relatively many parameters to fit the data. We addressed this question with both qualitative predictive checks and quantitative model comparison techniques, testing whether both dynamic noise and different priors for risky/safe options are strictly necessary to explain the empirical data^48,52^. To do so, we compared the PMC model to four simpler models. The first model was a baseline model where the noisiness of the first and second options were identical and the priors were the same for risky and safe options (model A^27,35^). Note that this model A is equivalent to a standard probit psychophysical model, which models the precision with which decision makers can compare two numerosities (with a potential bias towards one of the two options). In the second model, the noisiness of the first and second options was constant but the priors for risky and safe options could be different (model B). In the third model, the noisiness of the first and second options could be different but the prior for the risky and safe options was shared (model C). Finally, we also compared our models to a standard expected utility model (Model D^4,53^). The posterior predictive plots are presented in Figure 3. Visually, it is evident that none of the alternative models can explain the interaction effect of magnitude and order on the proportion of risky choices. Moreover, a formal model comparison using the Expected Posterior Log Density^48^ shows that the PMC model is vastly superior to the other four models. The difference in ELPDs is at least 21 standard errors (Figure 3E). When we used model stacking^54^, the model weight of the PMC model was 95%. These measures thus provide evidence for both mechanisms (i.e., varying noise for order and varying priors for riskiness) that are incorporated in our model. To make sure that our model comparison approach was sufficiently powerful, we also simulated 100 data sets using the 5 different models and used the same technique to estimate which was the generating model. When the PMCM was the generating model, it was correctly identified 100/100 times (See Supplementary Text 2 for more details). All in all, we demonstrate that the PMC model makes counterintuitive qualitative predictions that are fully in line with empirical data and that can only be captured by this particular model.

**Figure 3:**
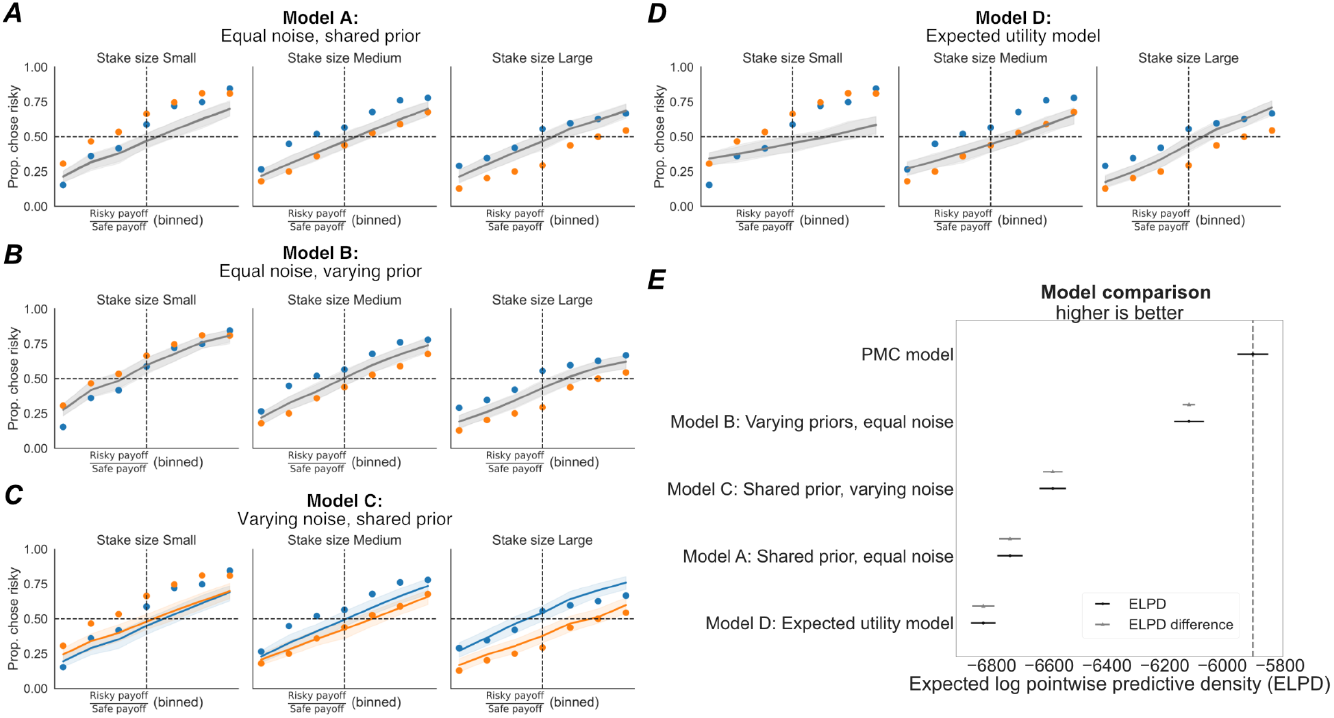
**A-D)** Posterior predictive plots for alternative models of different complexity. Unlike the PMCM, none of the tested models can explain all qualitative patterns in the data. Compare to Figure 2A. **E)** For formal model comparison, we used expected log-predictive density (ELPD), which estimates the likelihood of unseen data given the model parameters, making it an effective measure of out-of-sample performance^48^. ELPD comparison demonstrates that the PMC model significantly outperforms the other models tested (error bars are standard errors on the ELPD).

Finally, posterior estimates for the group mean of the model parameters (Fig. 2C) revealed that, as expected, participants had noisier payoff representations for first-presented versus the second-presented option (*p*_*Bayesian*_< 0.001 for both sessions). Notably, while at the group level, the subjective prior for risky payoffs did not have a higher mean than the prior for safe payoffs (3T: *p* _*Bayesian*_ =0.16, 7T: *p* _*Bayesian*_ =0.37, both: *p* _*Bayesian*_ =0.16), there were 13 participants who had a higher mean prior for risky than safe options (95% credible interval of the difference does not overlap with 0, see Fig. 2D). These individual parameter estimates suggest that some participants took into account that risky payoffs are higher on average whereas others do not.

In sum, our novel experimental paradigm and the PMCM can mechanistically characterize how a common and relevant context effect (whether options are held in working memory or perceived immediately) impacts the risk attitude of the participants. The model can explain why the experimental population is risk-averse in some contexts but risk-seeking in others, and it can account for individual differences in risk attitudes.

### Decoded payoff representations in parietal cortex predict noisiness and bias in choice

Having established that individual risk attitudes and changes in those attitudes across different choice contexts can be explained by our Bayesian perceptual account of risky choice, we addressed the final question of our study: Where do the trial-to-trial fluctuations in risk attitudes come from? We hypothesized that some choice variability –over and above the variability determined by context and individual differences– can be explained by momentary fluctuations in the acuity of neural representations^55–58^. Our previous work has shown that individual differences in the acuity of parietal magnitude coding predict individual differences in risky choice behaviour: Participants with noisier payoff magnitude representations tend to resort more to their prior beliefs and are, on average, more risk-averse^35^. This paves the way for the question whether there is a comparable relationship between neural noise of choice problems representations and behaviour *within* single decision-makers (in addition to neural noise in putative downstream value computations^59^). Such a relation could in principle be expected based on findings in perceptual neuroscience: For examples, trial-to-trial fluctuations in judgments about the orientation of Gabor patches are predicted by fluctuations in orientation representation in primary visual cortex^40,41,60^, and judgments about movement directions relate to differences in neural activity in middle temporal area (MT)^61,62^.

To be able to test this hypothesis, we first fitted a numerical receptive field (nPRF) model^35,43^ to single-trial BOLD responses locked to the *first payoff* presentation, to prevent any decision-related activity from contaminating our measures of neurocognitive representation (Fig. 1A). The nPRF model assumes that patches of cortex are tuned to numerosity - i.e., they have a preferred magnitude to which BOLD responses are largest - and that the size of the response to a stimulus decreases exponentially with its logarithmic distance to the preferred numerosity. Numerically tuned cortical areas were similar to previous studies: The surface-based locations where the model predicted single trial responses best (high explained variance, *R*^2^) largely followed the parietal mask based on our earlier study^35^ (see Fig. 4A). Within this mask, vertex-wise *R*^2^ values correlated significantly across the 3T and 7T fMRI sessions within participants with, on average, r = 0.25 (t(29) = 6.2, p<0.001). The vertex-wise preferred numerosities correlated with r = 0.15 (t(29) = 6.0, p<0.001). These results confirm that the intraparietal sulcus contains participant-specific patterns of numerosity encoding that generalize across sessions and even MRI scanners.

**Figure 4:**
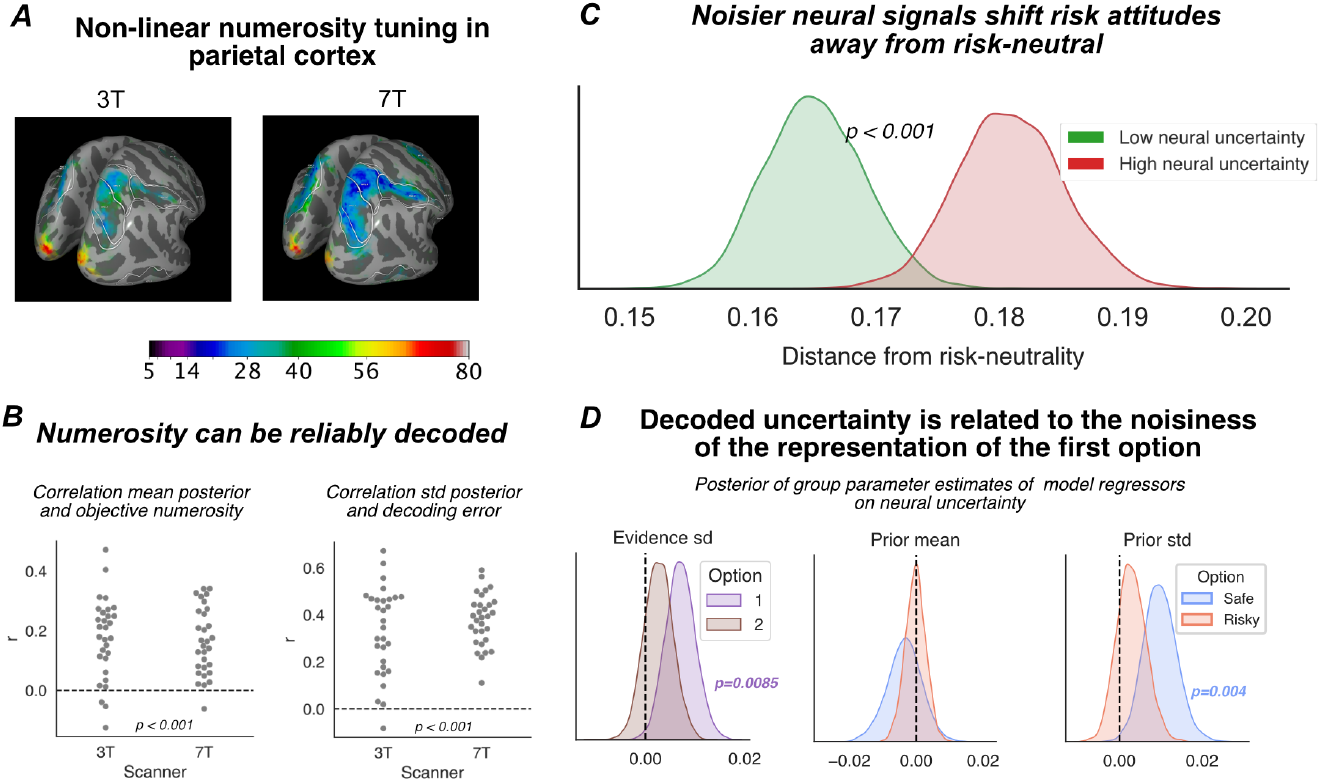
**A)** Numerical population receptive field (nPRF)-mapping reveals non-linearly tuned numerosity-sensitive BOLD responses around the intraparietal sulcus, closely following the ROI drawn on an independent data set^35^. Here we show the average preferred numerosity across all participants in fsaverage-space. **B)** Almost all participants (27 out 30 for 3T, 29 out 30 for 7T) show a positive correlation between the presented stimulus numerosities and decoded posterior means for single trials from out-of-sample fMRI data. Trial-to-trial fluctuations in the width of the decoded posterior are indeed predictive of the objective error of the mean posterior. **C)** Distribution of plausible values of the individual differences between risk-neutral behavior (RNP = 55%) and their actual indifference points. Thus, a distance of 0 would correspond to perfectly risk-neutral behavior. Choice behaviour on trials with a higher decoded neural uncertainty tends to be more biased: The RNP is further away from risk-neutrality (55%). The panel shows posterior estimates of the average participants-wise distance to risky-neutrality (abs(RNP - .55)). Thus, if participants were already risk-seeking on a particular trial type (particular order and particular safe payoff), they behaved even more risk-seeking when neural noise increased. The opposite effect is observed for participants and trial types for which behaviour was risk-averse. **D)** When the main six parameters of the PMCM were allowed to vary linearly with trial-to-trial fluctuations in the decoded neural uncertainty, both the noisiness of the representation of the first option and the dispersion of the prior on safe payoffs increase with neural uncertainty decoded from the first payoff stimulus. The noisiness of the second option, the location of the priors, and the dispersion of the prior on risky payoffs were not related to neural uncertainty.

After successfully fitting the nPRF model, we used a Bayesian inversion scheme to decode posterior estimates of presented numerosities from unseen fMRI data (leave-one-run-out cross-validation)^35^. Note that this method yields the posterior distributions over numerosity given the empirically observed single-trial brain activity pattern. Thus, the analysis does not only give us a point estimate of the numerosity expected to have elicited this BOLD pattern (mean of the posterior) but, crucially, also a measure of the uncertainty surrounding that point estimate. Earlier work in perceptual neuroscience has shown that this posterior uncertainty surrounding a stimulus feature decoded from brain activity predicts the randomness and bias of choices and judgments on those very stimulus features^35,40,41^.

Before linking neural uncertainty to behaviour, we first assessed the acuity of the decoder. The presented payoff magnitudes were decoded well above chance on the level of single trials, with an average correlation between actual numerosity and decoded numerosity of r=0.176 (3T; t(29) = 7.1; p<0.001) and r=0.16 (7T; t(29) = 8.0, p<0.001; see also Fig. 4B). Importantly, the decoded neural uncertainty (s.d. of the decoded posterior) was significantly correlated with the error of the decoder (absolute difference between mean posterior and actual numerosity) with r=0.33 (3T; t(29)=9.8, p<0.001) and r=0.38 (7T; t(29)=18.8, p<0.001). Since the uncertainty of the decoder relates directly to how well it can actually decode a given neural pattern, we can be confident that the variance in the decoded posteriors is meaningful and a useful proxy for the neural noisiness of the payoff magnitude representations that the decision-makers base their choices on^35,40,41^.

We also conceptually replicated an important result reported before by ourselves^35^ and others^63^, namely that participants with particularly noisy neural representations of magnitudes in the right parietal cortex also show noisier and biased behaviour. Specifically, participants who had a higher correlation between neurally decoded numerosities and the presented numerosities tended to have a less noisy representation of the first stimulus magnitude according to our behavioural PMCM (the correlation between decoding correlation and the dispersion of the likelihood of the first option, *v*_1_, r(29) =-0.48, p =0.007 for 3T and r(29) = −0.42, p=0.021 for 7T). Importantly, participants for which we decoded posteriors with, on average, a higher standard deviation also had a larger *v*_1_-parameter according to the PMCM (r(29) = 0.44, p=0.014 for 3T and r(29) = 0.38, p=0.039 in the 7T data), thus replicating our earlier findings^35^ twice, using different scanners and field strengths.

After reconfirming with these replications both the robustness of our IPS-signal decoder and the relevance of these parietal regions for individual risk attitudes, we proceeded to test whether *within each participant*, trial-to-trial fluctuations in apparent risk attitudes are related to latent fluctuations in the neural code that represents the potential payoffs that are at stake. To test this hypothesis carefully, we broke it down to two subhypotheses: On trials where neural payoff representations are particularly noisy, choice behaviour should (1) become less consistent and (2) more biased towards prior beliefs. In the PMCM, both effects can jointly be explained by an increase in noise. However, we aimed to first test both hypotheses separately. To do so, we used a standard psychophysical model that fits a psychophysical curve with both an indifference point and a psychometric slope to the choice data as a function of the log-ratio of the risky and safe payoffs^27,35^. To make sure we found an effect of neural uncertainty over and above any order- and magnitude-effects, we performed a median split separately for every possible first-presented payoff and fitted separate psychometric curves to these two sets of trials that were matched in payoffs and order but showed relatively high or low neural noise.

In line with our expectations, the psychophysical slope was substantially shallower for trials with a higher decoded neural uncertainty (*p* _*Bayesian*_ =0.0458 for the 3T data set; *p* _*Bayesian*_=0.0238 for the 7T data set). Thus, the noisier the signal was according to our neural decoder, the more randomly (i.e., independent from the presented payoffs) participants chose between the options. To test whether participants were more biased in their perception of the potential payoffs when their neural representations were particularly noisy, we tested whether estimated indifference points were further away from risk-neutrality when neural noise was higher. To quantify risk-neutrality, we again used the *risk-neutral probability* (RNP). We fitted a hierarchical Bayesian psychometric model and sampled the resulting participant-, payoff- and order-specific RNPs. These RNPs were indeed farther away from risk-neutrality (55%) when the neural uncertainty was higher (18.5 percent points, s.d. 9.2 versus 16.7 percent point, s.d. 9.2;*p* _*Bayesian*_ <0.001, Fig. 4C; these results replicate when separately estimated for two sessions, *p* _*Bayesian*_ = 0.012 for 3T and *p* _*Bayesian*_ =0.013 for 7T). Thus, the less precise the numerosity-tuned responses in IPS according to our decoder, the more participants reverted to their prior beliefs and the less risk-neutral they became in their risk attitudes.

Finally, as the ultimate test of the hypothesis that trial-to-trial fluctuations in risk attitudes are (partly) driven by fluctuations in neural noise, we tested whether an increase in the noise parameter of the PMCM can explain the observed link between neural acuity and behavioural consistency and choice. We hypothesized that, in terms of PMCM parameters, the first choice option should be represented more noisily when the corresponding decoded signal was less precise. Therefore, we refitted the PMCM to behavioural data, but now all six model parameters were estimated as a linear sum of a (participants-wise) intercept and a regression parameter that was multiplied with the z-scored trialwise neural uncertainty measures (i.e., the standard deviation of the decoded posterior). If this latter regression parameter differed from 0, the corresponding latent variable linearly varied with neural uncertainty^64,65^. This analysis confirmed that both the noisiness of the first option (3T: *p* _*Bayesian*_ =0.02; 7T:*p* _*Bayesian*_ =0.04; both: *p* _*Bayesian*_ =0.009; See Fig. 4 and Supplementary Fig. 3) and the standard deviation of the prior on safe payoffs (3T: *p* _*Bayesian*_ =0.02, 7T, *p* _*Bayesian*_ =0.01; both: *p* _*Bayesian*_ =0.004; See Fig. 4 and Supplementary Fig. 3) linearly increased with increased neural uncertainty. These results align with the psychophysical modelling results: When neural noise is high, the representation of the payoff of the first option becomes noisier, resulting in larger CT biases and more variable behaviour. However, the increased dispersion of the prior belief about safe payoff magnitudes with increased uncertainty reveals that it is mostly the risky (rather than safe) options that drive the increased CT bias with increased neural uncertainty.

In conclusion, we successfully decoded payoff magnitude representations during economic choice from right parietal cortex and replicated earlier findings of a link between the acuity of an individual’s numerical magnitude representation in the IPS and their choice consistency and risk attitudes. Crucially, we now found that even trial-to-trial fluctuations in the acuity of payoff representations in parietal cortex were mechanistically linked to trial-to-trial fluctuations in choice consistency and risk attitude. This provides direct biological evidence that the endogenously fluctuating precision of neural magnitude representations underlies time-variations in revealed risk attitude.

### Choices in symbolic presentation format also reveal shifts in risk attitude as a result of order and stake size

In the fMRI experiment, we used dot clouds to present potential payoffs, because such payoff stimuli gave us the most reliable signal to probe neurocognitive representations in the parietal lobe using fMRI^35,43^. One might wonder, however, if the observed effects of presentation order and their interaction with stake size might be specific to this non-symbolic presentation format of dot clouds. The non-symbolic presentation format we used may also introduce some stimulus ambiguity that might be weaker in more traditional economic choice tasks using Arab numerals (although there can be profound noise and bias in the perception of symbolic numerical magnitudes as well^37,66,67^). To address these concerns, we conducted an additional experiments with 58 participants (average age 24.6, range 20-41; 31 females) who performed a very similar choice task, but now the dot clouds were replaced with Arab numerals (e.g., “CHF 20,10” or “CHF 10.08”; see Fig. 5A).

**Figure 5:**
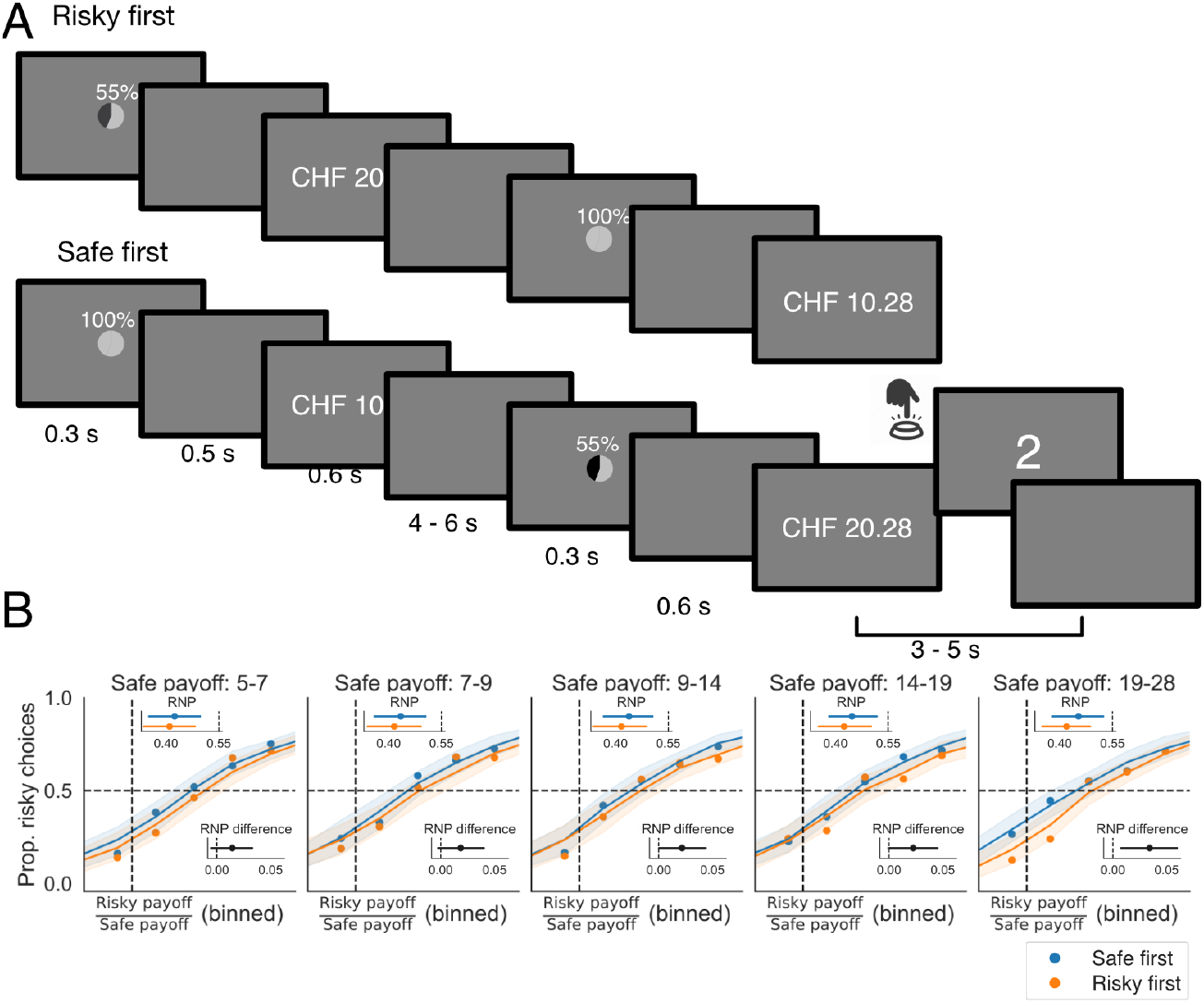
**A)** Additional experimental paradigm that uses Arab numerals instead of dot clouds to present potential payoffs. **B)** Average psychophysical response curves for different stake sizes, estimated using a hierarchical psychophysical model. The x-axis represents the ratio between the risky and safe option (log-space), and the y-axis shows the average proportion of risky choices. Raw data are plotted as dots, with the shaded area indicating the 95% credible interval of the posterior predictions from the psychophysical probit model. The dashed black line denotes the indifference point for a strictly risk-neutral decision-maker (1/0.55). RNP (Risk-Neutral Probability) values for each condition are provided for each stake size, along with the RNP difference between conditions. Participants exhibited greater risk-averse behavior when risky options were presented first, an effect that increased with stake size.

Just as for the fMRI data, we fitted a set of psychophysical choice models, where the probability of choosing the risky option is a (probit) function of the ratio of the risky and safe option. Choice consistency and risk attitude were modeled separately for different stake sizes. Formal model comparison again identified as the best model a full interaction model in which the indifference point (i.e., the risk neutral-probability, RNP) corresponded to even more risk aversion when risky options are presented first *and* they are relatively large (ELPD: −5383.73 for *order x stake* model vs −5430.28 for *stake* model, dSE: 11.09; see Figure 5B). Thus, this additional analysis is consistent with the behavioral results from the fMRI study: Participants become more risk-averse when risky options are presented first (and are therefore represented more noisily), in particular when stake sizes are large (and the noisy magnitude representations are biased more towards the smaller prior). This is also consistent with the interpretation of the CT account that risky options are underestimated relatively more when they are objectively large. Additionally, our results also replicate earlier findings^27,35^ that individual differences in choice consistency and indifference point (risk attitude) are significantly correlated (r(57) = 0.34, p=0.00822). This is again in line with the central assumption of the perceptual account of risk attitudes: Participants whose choices are based on noisier representations (and therefore more inconsistent) also deviate more strongly from risk-neutral behavior.

To account for the results more mechanistically, we also fitted the four different versions of the PMCM, with varying or shared priors for risky and safe payoffs, as well as different or equal amounts of noise for the 1st and 2nd presented option. Formal model comparison revealed that, just as in the fMRI study, only the full PMCM model could explain the empirical data and vastly outperformed a standard expected utility model; see Supplementary text 3 for more details. Ten out of the 58 participants robustly showed more noise for the 1st compared to the 2nd option, in line with working memory noise on the first-presented option. However, surprisingly, a smaller number of participants (5/58) robustly (*p* _*Bayesian*_ <0.05) showed more noise for the 2nd option than the 1st option, contrary to a memory noise effect. We believe these results might reflect differences in attentional strategies^68^, which could be investigated in future modeling work using eye tracking and/or choice-specific noise manipulations.

Taking stock, the key finding of this additional experiment is that the order in which options are presented shapes risk attitudes, and this effect interacts with overall stake size—even when using more traditional, symbolic stimuli to convey payoff magnitude. These results show that presentation order effects—and their interaction with stake size—are not specific to dot cloud displays but also generalize to more natural presentation formats. Moreover, the observed data using Arab numerals are well explained by our PMC model, whereas earlier perceptual models of risky choice and standard expected utility frameworks fail to capture the full pattern of results. This provides strong evidence—consistent with earlier work^27,35^—that perceptual processes and Bayesian inference play a key role in risky choice, even when information is presented in seemingly unambiguous, symbolic form. See Supplementary Text 3 for more details on this additional experiment.

## Discussion

Overt risk attitudes –and the latent underlying risk preferences – play a central role in many aspects of our behaviour, but we still don’t completely understand where they come from, and why they change across different contexts or even repetitions of identical choices. Here, we provide modelling and neurobiological evidence that rapid changes in individual risk attitudes can originate in the neurocognitive processes that underlie our perception of (monetary payoff) magnitudes. We introduced a new experimental paradigm that modulates the noisiness with which different choice options are perceived, by presenting them in different orders, as is typical for everyday choice situations in which options are considered one after another. This sequential presentation made choice options more/less prone to working memory noise^45,46^, resulting in central tendency effects^24,34^ that led participants to perceive the very same choice options as larger or smaller depending on their presentation order, magnitude, and riskiness (reflected in different priors for risky and safe options). We captured these effects with a new Bayesian model of risky choice - the PMCM - that explicitly accounts for dynamic fluctuations in the relative noisiness with which different choice options are perceived, as well as for different prior expectations based on the context of a particular choice problem. Moreover, by using fMRI and inverted encoding models, we could provide neural evidence for such rapid fluctuations in neurocognitive noise, showing that these predict risky choice in a manner consistent with the PMCM.

Our results have profound implications for both theoretical and empirical work on risk attitudes. First, in line with other recent work^13,14,17^, they defy the conventional economic wisdom that risk attitudes are stable^2^ and demonstrate that they can systematically vary due to fluctuations in the noisiness of neurocognitive representations of the relevant decision variables. This relevance of noise in the decision-making process for risk attitudes dovetails with earlier work highlighting that differences in risk attitude may easily be conflated with differences in choice consistency^47,69,70^. However, rather than assuming that noise may affect choices unsystematically, at a late stage of the decision process, our model emphasizes that noise originates early in the decision process, during perception of the choice-relevant information, thereby determining risk attitudes already before any actions are taken. These perceptual influences on overt behavior pose challenges to the common approach of interpreting choices as revealing (only) latent risk preferences.

A second key implication of our study – congruent with other recent work^27,71,72^ – is that a large part of inter- and intra-individual variability in risky choice behaviour can be explained by fluctuations in basic perceptual neurocognitive representations, over and above any possible differences in subjective valuation of the choice options. We show that decision-makers can seemingly reverse their risk attitude (i.e., become risk-seeking rather than risk-averse) simply due to changes in the order in which choice options are presented. This raises the crucial question to what extent variability in risky choices really reflects subjective valuation (rather than noisy perception), as usually assumed in standard economic models. Notably, our PMC model, which contains no subjective valuation process, could explain a wide range of choice patterns across subjects and contexts, whereas a standard economic choice model based exclusively on subjective evaluation (the EU model) failed to do so. While this evidently does not prove that subjective valuation is not involved in risky choices, it still suggests that perception may play an important role and needs to be accounted for, at least in the small-stake laboratory experiments widely used to study risk preferences.

This substantial impact of perception on risk attitudes (rather than valuation) might explain the ‘risk elicitation puzzle’: the fact that risk attitudes (or “elicited preferences”) are only marginally correlated across elicitation methods^6^. Moreover, the central role of perceptual effects in risk attitudes may have profound implications for welfare economics, which often relies on elicited preferences to calibrate models of the impact of economic policies^73^. Thus, future work should attempt to further unravel potentially separate contributions of neurocognitive processes related to subjective valuation versus perception to individual preferences and, if possible, develop economic choice tasks where perception can be dissociated from participant’s subjective valuation.

Future studies attempting to dissociate valuation and perception should systematically manipulate both sets of processes. Valuation may be experimentally manipulated by changing objective choice properties and comparing qualitatively different decision-makers, while perception may be influenced by orthogonally manipulating the acuity of the choice-relevant information. Here we did so by representing potential payoffs with non-symbolic stimulus arrays and changing their order.

One might argue that this choice context may induce effects in a purely ‘low-level perceptual’ manner that should be absent for choices with more common Arabic numerals. However, there is ample evidence that the perception of Arabic numerals is prone to similar perceptual effects^67^. For example, when participants have to make decisions about rapidly-presented Arabic numerals, they are sensitive to order effects^66^ and the underlying statistics of the presented numerals^74^, just as for the non-symbolic presentation formats in the study presented here. Furthermore, recent neuroscience studies have shown that numerosity-tuned regions similarly represent both symbolic Arabic numerals and non-symbolic stimulus arrays^75^. In line with this, our recent work^35^ has shown that the performance on a perceptual task involving the numerosity of stimulus arrays similarly predicts choice consistency and risk attitude in risky choice tasks that present payoffs either non-symbolically or as Arab numerals. Finally, in the present study, we also presented a choice task where the non-symbolic stimulus arrays were replaced with unambiguous, Arab numerals. In that experiment, we also found an effect of order on risk attitude, as well as an interaction with stake size. A closer look at the behavior and parameter estimates of individual participants showed a wide array of potential choice behavior that could not be explained by a standard expected utility model (which was also confirmed with formal model comparison measures). All this highlights that choice problems presented in non-symbolic and symbolic presentation formats engage similar neural magnitude representations, and that our model can capture common processes underlying choices about payoffs presented in both presentation formats..

The findings that our Bayesian model of the inference underlying perception could account for several counterintuitive context effects on risk attitudes, for choice sets both ambiguous and non-ambiguous payoff stimuli, suggests that this type of model may be important for understanding several other effects documented in the literature. For example, another well-established determinant of the acuity of choice information is the time that choice information is attended to. Indeed, eye-tracking data show that both the order and duration with which participants attend to relevant choice variables profoundly influence economic choice^76^. Importantly, participants act *as if* they multiply relative values of choice options with the amount of time they gaze upon a choice option^77,78^. This multiplicative effect is fully consistent with a Bayesian account where attention increases the acuity with which we perceive information. Thus, particularly valuable options are chosen more often with increased attention, but less-valuable options less often (since the CT effect will decrease as a function of dwell time)^79^. Another way in which perceptual processes preceding putative valuation may affect risk attitudes are changes in the statistics of the local environment^80^. Indeed, it is well-known that economic behavior is sensitive to the larger choice set across an experiment^14^. Future studies should thus attempt to further test with rigorous Bayesian modelling whether these choice set effects are not only qualitatively but also quantitatively consistent with a model where subjective priors adapt to the local choice context^24,30,81^.

Our work also has implications for existing neuroeconomic studies and opens up some exciting new research directions. First, studies attempting to identify neural correlates of subjective value^82^ need to take into account that at least under some conditions, subjective values inferred from choice behaviour might be a compound measure that also includes perceptual noise and biases, which can profoundly influence any decision variable constructed from this information. This means that neural correlates of subjective value might include perceptual areas as well as ‘true’ valuation areas^83,84^. However, our work also suggests that computational modelling of behavioural and neural data could help to decompose these sources of choice variability^85^. It will be interesting to see whether explicitly incorporating perception in such models can help to elucidate whether subjective value is represented in the brain completely independently from perceptual variables.

Second, our results suggest a key role for parietal regions in economic decisions that involve (numerical) magnitudes. Most neuroeconomic theories propose that, during choice, subjective values are represented in the ventromedial and/or orbitofrontal cortex (vmPFC/OFC)^86^, whereas parietal cortex is generally conceptualized as a brain region that integrates the final evidence for different choice options based on subjective or objective stimulus properties^87–89^. However, our results suggest that, at least when magnitudes are involved, parietal cortex also directly represents initially choice-relevant stimulus information during perception of the choice problem. This provides a new explanation for earlier discoveries of neural substrates of subjective value of specific choice options in the parietal cortex of non-human primates^90,91^, as well as recent work using brain stimulation that shows that activity in parietal cortex is causally involved in risky choice^92,93^.

Clearly, the precise role of parietal cortex in economic choice remains to be fully elucidated and is likely to depend on the specifics of the choice task, as well as the phase of decision-making during which one monitors parietal activity. However, one promising working hypothesis could be that when objective magnitudes are involved, parietal cortex can both represent relevant magnitudes (e.g., probability of payoff^94^ or monetary amounts^35^) of the option currently under consideration/attention^94,95^, as well as accumulate evidence for choice options as also found in recent work on perception^96^. Notably, almost all work in (human) neuroeconomics has assumed linear coding of subjective value in large cortical locations that adhere to macro-anatomy (i.e., MNI space). However, our work suggests a non-linear code that is only loosely related to macro-anatomy – at least for parietal representations of potential payoffs (see also work on non-linear coding of probabilities^94^). More sophisticated fMRI analysis approaches may thus be needed to fully delineate representations of objective stimulus information and subjective values during economic choices^84^. For instance, studies decomposing subjective value in its constituent parts (basic perceptual and subjective) should aim to use analysis frameworks that allow for non-linear codes and take into account individual functional neuroanatomy. An exciting open question for such work may be how fluctuations in early perceptual representations of a choice problem (e.g., potential payoff) covary with prefrontal representations of subjective value.

A central mechanism in Bayesian theories of (economic) cognition are prior beliefs about the state of the world. Some have argued that this may make Bayesian models of cognition overly flexible^97^ (but see^98^). In the PMCM, decision-makers have prior beliefs over the plausible ranges of risky and safe payoffs. Because the mean and spread of these priors are free parameters, the PMCM is indeed a relatively flexible model. However, both formal model comparison and careful qualitative inspection of the empirical data show that less flexible models of risky choice (both of perceptual and preferential nature) can simply not account for the qualitative behavioral patterns in our data. Having said that, future work should validate whether change in the objective distributions of payoffs will result in the predicted behavioral changes and corresponding parameter shifts of the PMCM. Encouragingly, earlier work on both risk^30^ and loss aversion^14^ has already shown that human decision-makers are sensitive to the statistical properties of earlier choice problems, in line with Bayesian theories of economic choice. An important, related question is how, and on which time scale, human decision-makers can learn and adapt their subjective priors. After some initial attempts at including such a learning mechanism in our model, we believe that the choice paradigm we employed here is not suitable for such questions, because the amount of information present in a single accept/reject decision is limited. Therefore, future work might employ different paradigms with a more continuous behavioral measure, such as willingness-to-pay^99^.

A related critique of Bayesian theories, and potentially the PMCM, is that they are not mechanistic^97^ models of neural function but rather prescriptive models of information processing. Nevertheless, the PMCM does give us an increased understanding of the mechanism by which risk attitudes arise, through the lens of Bayesian perceptual inference and its link to neural activity. Concretely, without an understanding of working memory noise and the formalisation of its effects using the Bayesian framework, it remains very challenging to explain the rather complex observed stake x order interaction. Moreover, our neurocomputational modeling results link the abstract Bayesian formalism to concrete, neural signals that we observe using neuroimaging: Payoff magnitudes are represented in a non-linearly tuned population code in parietal cortex, which correlates with a posterior belief. The less consistent this population code across its subpopulations, the more uncertain the decision-maker’s beliefs should be^32^ and the more they should revert to their prior beliefs. Exactly this link between neural and behavioral signals, which forms a concrete decision-making mechanism, is what we observed in our data. While much work remains, our findings show how abstract, formal Bayesian theories of cognition can be integrated with measurements of corresponding neural population codes, which offers an interesting approach for gaining a more mechanistic understanding of decision-making.

Our work aligns with a recent body of research that conceptualizes economic choices as rational responses to cognitive capacity constraints^16,25,71,100,101^ and makes three new contributions to this literature. First, we present a computational model that can be used to estimate how choice contexts modulate both prior beliefs and perceptual noise associated with different choice options. Second, our PMC model can mechanistically explain how discrete categories of risk attitudes (i.e., risk-averse, risk-neutral, risk-seeking) can be understood as a continuum emerging from the interplay between subjective beliefs about potential payoffs and noisiness of perception and can predict corresponding changes in risk attitude across different contexts. Third, we present neuroimaging evidence that fully endogenous fluctuations in the noisiness of *neural* representations of economic variables^53^ determine risk attitude in line with a Bayesian perceptual account. This last finding also highlights the promise of using advanced neuroimaging data and analysis methods to help validate and refine computational models of economic choice^102^. Future work in this direction may, for example, link risk attitudes to the noisiness of neural representations of other relevant economic choice variables, like probability^71,94,99^, time^103,104^, and social constellations^105,106^.

Finally, our results also have potential clinical and policy implications. First, the abundant preference for gambling has long been a puzzle for economists^107,108^. Might the preference for gambling be explained as a misperception of the potential payoffs? One relevant finding in pathological gambling is increased arousal during gambling situations^109,110^, which is known to interact with perceptual biases^111^. Indeed, gamblers report unrealistically optimistic beliefs about potential outcomes^112–114^. Future work might investigate false beliefs in pathological gamblers using explicit Bayesian models of risk perception^115,116^. Such false beliefs might have important policy and moral implications for the public communication about gambling. Our results also have implications for the domain of finance. In personal banking, it has become common practice to elicit risk preferences when taking on new clients^117^. If perceptual effects partly drive such elicited preferences, these estimations might be erroneous and clients might be offered an investment portfolio that is overly defensive, just because they have relatively noisy numerical perception or overly pessimistic beliefs about potential long-term stock returns.

To conclude, our work reveals that perceptual biases to presentation order and relative magnitudes, as well as brain state, can profoundly impact an individual’s risk attitude, and they introduce an experimental and modelling approach that can be used to systematically investigate such effects for a whole range of economic choices. Our findings empirically substantiate emerging proposals that economic choice behavior should be understood as the fluctuating output of a capacity-limited brain rather than fixed as-is economic preferences^16,25,27,100,101^. Our new model and experimental approach is highly relevant to (neuro)economic research because it formalizes why there might be no such thing as an ‘objective’ risk attitude that is completely independent of context and brain state^6,118^, and because it allows researchers to quantify the corresponding effects. More generally, our study demonstrates that attempts to elicit (risk) preferences need to consider the substantial variability in responses and should be mindful of potential confounds like presentation (or fixation) order and the general range of options offered during an experiment. These considerations can profoundly impact real-world practice as well, for example when risk elicitation paradigms are increasingly used to compile investment portfolios^117^ or characterize the symptoms of psychiatric disorders^119,120^.

## Methods

### Participants and ethics

Thirty right-handed participants (14 females, ages 20 to 34) volunteered to participate in this study. We informed them about the study’s objectives, the equipment used in the experiment, the data recorded and obtained from them, the tasks involved, and their expected payoffs. We also screened participants for MR compatibility prior to their participation in the study. No participant had indications of psychiatric or neurological disorders or needed visual correction. Our experiments conformed to the Declaration of Helsinki, and our protocol was approved by the Canton of Zurich’s Ethics Committee.

### Procedure

Participants were invited for a total of 4 sessions. Two sessions took place at the 3T scanner and two sessions took place at the 7T scanner (see ‘MRI parameters’ for more information on scanner type and scanning parameters). Participants always first performed two sessions at one scanner, before the other two at the other scanner. Seventeen (17) participants performed the first two sessions at the 7T scanner, the other 13 at the 3T scanner. During the first and third session –so both at the 3T and 7T scanner– participants performed a calibration version of the risky choice task so we could estimate their average risk attitude and choice consistency within a specific scanner environment before they performed the main task of interest. The calibration task was done in the scanner while undergoing anatomical scans. Participants made choices about 96 possible risky gambles (the safe payoff was either 5, 7, 10, 14, 20, or 28; the risky option was a 2^*h*/4^ times the safe option with h being all integers between 1 and 8). These offers were presented twice, once with the safe option first and once with the risky option first. A psychometric probit model with “chose risky option” as the dependent variable and an intercept and the log-ratio of the risky and safe option ^27^ as independent variables was fitted to these calibration data. The fitted model established the precise mapping between risky/safe payoff ratios and the proportion of risky choices. We then made a participants-tailored design with 8 fractions equally spaced in log-space that –according to the psychometric model– were predicted (equally-spaced) proportions of risky choices between 20% and 80%. These 8 fractions were combined with 6 safe payoffs (5/7/10/14/20/28). Also, all these offers were presented twice with the safe option first and twice with the risky option presented first (so, for every session, each possible safe/risky/order-permutation occurred twice). This amounted to 192 trials per session. After the anatomical scans and the calibration task, participants performed a numerosity mapper task following Harvey et al. (2013; data not shown in this paper).

For the second session in a scanner, participants filled in all information and consent forms and then immediately performed 192 trials of the risk task in the scanner, spread out over 8 blocks of approximately 5 minutes each.

After all 4 sessions, we selected a random risky choice trial from each of the two calibration sessions and one of each of the main task sessions. For the risky trials, a digital random number generator was used to hand out the risky option 55% of the time. The participants got the average payout across the 4 sessions on top of their hourly fee (30 CHF per hour; approximately 30-35 USD).

### Experimental paradigm

The task of the participants was, for every trial, to choose between a certain amount of money (5/7/10/14/20 or 28 CHF) or a gamble with 55% probability of winning a larger amount of money and a 45% probability of winning nothing at all. The choices were presented by a sequence of tailored stimuli. The general sequence is illustrated in Fig. 1A. The screen always contained a red cross with two diagonal lines to keep fixation near the centre of the screen and to not confound the numerosity of stimuli like a standard fixation cross or point might^43^. The start of a new trial was indicated by the fixation cross turning green for 250 ms. Then, after a pause of 300 ms with just a red fixation cross, a pile chart with a diameter of 1 degree-of-visual-angle (dova) was presented for 300 ms to indicate the probability-of-payout for the coming stimulus. This was always either 55% or 100%. Then, after another 500 ms of just fixation cross, a stimulus array of 1-CHF coins appeared that represented the potential payoff of the first choice option. The coins had a radius of 0.3 dova (degrees of visual angle) and were all randomly positioned with their centre within a circular aperture with a diameter of 5.25 dova. The coin stimulus array was presented for 600 ms, after which only the fixation cross was presented for a jittered duration of either 5, 6, 7, or 8 seconds. Then, a piechart indicating the probability of the second payoff was presented for 300 ms, followed by a 300ms fixation screen, and another coin stimulus array, representing the potential payoff of the second option, again for 600 ms. As soon as the second stimulus array was presented, participants could indicate their response with their index (first-presented option) or middle finger (second-presented option). As soon as they responded, they saw a 1 or 2 indicating which option they had chosen for 500 ms. After the coin stimulus pile was presented, the remaining duration of the trial was either 4, 4.5, 5, or 5.5 seconds. During the time that was left after the feedback stimulus, participants had to indicate “how certain” they were about their choice using a 4-step Likert scale (by pressing response buttons with four fingers, from index finger-pinky). The results from this certainty rating are not presented in this paper.

dAt the 7T scanner, stimuli were presented using the Avotec (Avotec, FL, USA) binocular goggle system (800 x 600) with a field-of-view of approximately 25 dova.

## Behavioural analyses

### Psychometric/KLW model

#### Model specification

Khaw, Li, and Woodford^27^ showed that in risky choice paradigms where participants choose between a certain payoff *C* and a risky payoff *X* with a probability of payout *p*, observed choice behaviour can be understood as the outcome of a Bayesian inference process.

Specifically, their model (from on referred to as KLW model) assumes that participants do not make decisions based on the objective payoffs *C* and *X*, but based on noisy representations of those payoffs, described as random variables *r*_*x*_ and *r*_*c*_. The likelihood function describes the probability of noisy representations conditional on *C* and *X*:

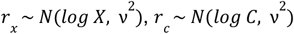

As can be seen in the formula, these representations take place in a logarhitmic payoff space. This means that participants will adhere to Weber’s law. Thus, in natural space, the randomness of their choice behaviour is predicted by ratios between options rather than absolute differences in natural space. Moreover, the parameter *ν* determines the noisiness of representations and thereby how variable behaviour will ultimately be.

Crucially, the KLW model also assumes that people apply a *common prior* for both the risky and safe options:

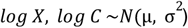

In the KLW model, the µ-parameter has no influence on predicted behaviour. However, because the decision-maker applies the same common prior to both risky and safe options, risky options will always *be relatively underestimated compared to the safe option* (this is because risky options always have higher payoffs than safe options for them to be attractive to the participants). The extent to which underestimation of larger payoffs happens is a function of the ratio between the variance of the likelihood *ν* ^2^ and the dispersion of the prior σ ^2^, β:

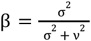

Specifically, the conditional expected values of the two options conditional on *r* are:

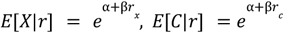

where α is a function of the mean and standard deviation of the prior:

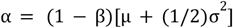

(Rational) participants will choose the risky option if and only if *p* ×*E*[*X*|*r*] > *E*[*C*|*r*] or, equivalently, log *p* + β*r* _*x*_> β*r*_*c*_, or log *p* + β*r*_*x*_− β*r*_*c*_ > 0 Since both *r*_*x*_ and *r*_*c*_ are (independent) Gaussian random variables, conditional on *X* and *C*, their difference is a Gaussian variable as well:

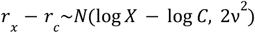

Crucially, this means that the probability that the participant chooses the risky option ( log *p* + β*r*_*x*_ − β*r*_*c*_ > 0) is conditional on *C* and *X* can be described using the cumulative normal distribution Φ(*x*):

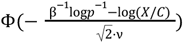

The likelihood of the KLW model is equivalent to that of a Generalized Linear Model (GLM) with the Normal CDF Φ as the link function and a Bernoulli error distribution (probit model). Such a GLM can be fitted using of-the-shelf functions in most standard statistical modelling software. When using a GLM-approach, the choice of the risky option is the dependent variable and an intercept (all 1’s) regressor and log(*X*/*C*) as a regressor. Crucially, the resulting intercept parameter δ and slope parameter γ from the probit model can then be re-expressed in the parameters of the KLW model:

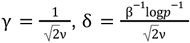

#### Parameter estimation

We fitted the psychometric model using the Bayesian hierarchical GLM-estimation package Bambi^121^. The standard probit model assumes that choices are determined by a fixed effect of both an intercept (determining the risk aversion) and the log ratio of the risky and safe option. We also estimate participants-wise random effects on all parameters so we had participants-wise KLW parameters:

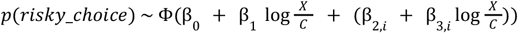

Where β_0_ and β_1_ are fixed effects and β_2,*i*_ and β_3,*i*_ are random (participant-specific) effects.

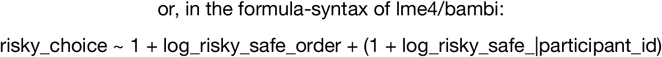

For some analyses in which we wanted to see how risk attitudes are modulated for different contexts, we also included the order and/or the value of the safe option in the model as independent variables, so:

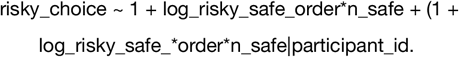

Note that the point-of-indifference of the probit model is a function of both the intercept δ and slope parameter 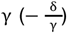. To meaningfully quantify the point-of-indifference, we used the risk-neutral-probability (RNP)^27,35^. This is the hypothetical probability of the risky option for which a risk-neutral decision-maker would show the same point-of-indifference as the participants. Thus, any RNP above 55% implies risk-seeking, whereas any RNP below 55% implies risk aversion. We defined participants as being risk-averse when the 95% Credible Interval of their RNP was below .55, and risk-seeking if the CI was above .55. When the CI overlapped with .55, we categorised participants as risk-neutral.

The risk-neutral probability can be derived from the slope γ and intercept δ as follows:

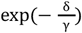

To fit the hierarchical probit/KLW model, we used the standard weakly informative priors as implemented by default in the Bambi (based on the approach of the well-known *rstanarm*-package) package^121^. Briefly, the priors are Gaussian distributions centred at 0, with as a standard deviation 2.5 times the ratio between the standard deviation of the dependent and independent variable^121^. To sample from the parameter posteriors, we used the no U-Turn sampler (NUTS), which is a self-tuning version of the Hammiltonian MCMC sampler and is particularly efficient for high-dimensional models with correlations between the parameter, such as our model here^122^. We collected 4 chains of 2000 samples each (1000 burnin). We always visually inspected the traces of the NUTS sampler and made sure the Gelman-Rubin statistic was below 1.05 for all parameters.

### PMC model

#### Model specification

Our Perceptual Memory-based Risky Choice model (PMCM) assumes that participants base their risky choices on noisy representations of the payoffs and choose the one with the largest expected payoff on a particular trial. Crucially, in the PMCM, the noisiness of the representation may depend on the order of presentation:

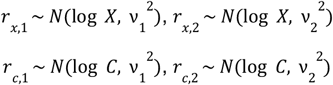

Where *r*_*x*,1_ pertains to the representation of the payoff of the risky option presented first, *r*_*x*,2_ to the representation of the payoff of a risky option presented second, *r*_*c*,1_ to the representation of the payoff of a safe option presented first, and *r*_*c*,2_to the payoff of a safe option presented second. Note that the noisiness of the risky and safe options *ν*_*i*_ is the same, only order of presentation influence noisiness. We expected that the noise for the first option *ν*_1_ will be higher than that of the second option *ν*_2_, but this is not enforced anywhere in the model (e.g., they have identical priors in the estimation procedure).

The PMCM also assumes that participants (potentially) employ different priors for the risky and safe payoffs, because these payoffs come from objectively different distributions:

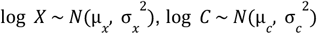

For a given parameter set [*ν*_1_, *ν*_2_, µ_*x*_, µ_*c*_, σ_*x*_, σ_*c*_] and objective payoffs *X* and *C*, we can now obtain the distribution of the expectations of the participants for the two options, which is the product of the prior and evidence distributions and follows a normal distribution (in logarithmic space) :

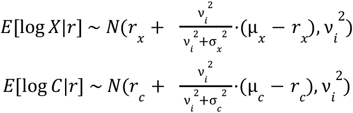

Thus, the distribution of differences between these expectations in log space is given by the difference of these two distributions, *E*[log *X*|*r*] - *E*[log *C*|*r*], which is also a normal distribution:

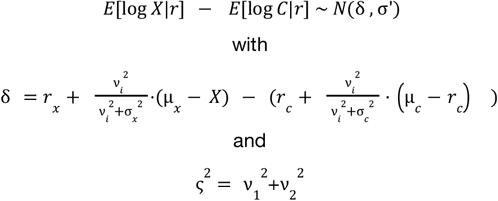

The likelihood of choosing the risky option according to the PMCM is defined as the probability that the participant’s estimate of the difference between the risky and safe option ( *E*[log *X*|*r*] − *E*[log *C*|*r*]) is larger than the payout probability of the risky option in log space:

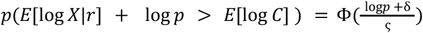

Where Φ(*x*) is the standard cumulative normal distribution.

#### Model estimation

We implemented the PMCM using the Bayesian statistical modelling library pymc (v.5)^123^ and wrapped it in our Python package *bauer*^*1*^. We used a hierarchical (regression) approach for estimation. This means that for all main six parameters of the model [*ν*_1_, *ν*_2_, µ_*x*_, µ_*c*_, σ_*x*_, σ_*c*_], we assume a group distribution which is a Gaussian distribution. Furthermore, for the *ν*- and σ -parameters, which are necessarily non-negative, we estimated transformed parameters in an unrestricted space [− ∞, ∞] which we then transformed into the non-negative domain [0, ∞] using the softplus function log(1 + exp(*x*)).

For a given parameter θ (e.g., *ν*_1_ or σ_*x*_), the participants-specific parameter of participants *p* was a linear combination of the mean group parameter µ_θ_ plus an individually determined offset-parameter δ_θ_ times the standard deviation of the group distribution σ_θ_ :

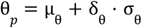

This offset-based specification of the hierarchical model was chosen to prevent the likelihood “funnels” that plague high-dimensional hierarchical models^124^.

When relating the PMCM parameters to neural measures, we used a regression approach^64^, where a given parameter θ_*p*_ (e.g., *ν*_1_ or σ_*x*_) of participants *p* on a trial *t* was a linear sum of both an intercept θ_0_ and a regression coefficient θ_*n*_ times the neural measure at trial *t, n*_*t*_ :

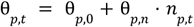

We used mildly informative priors on all group distribution parameters:

For the evidence parameters:

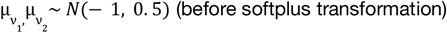

For the mean of the prior on risky payoffs:

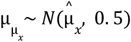

Where 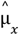 is the *objective* mean of the payoffs of risky options (in log space)

For the mean of the prior on safe payoffs:

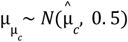

Where 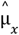 is the *objective* mean of the payoffs of safe options (in log space)

For the standard deviation of the priors:

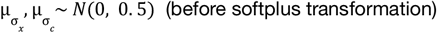

The regression coefficient on (z-scored) neural measures was a Gaussian centred at 0 with a standard deviation of 1:

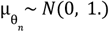

The prior on all group variance parameters was a Half-Cauchy parameter ^125^:

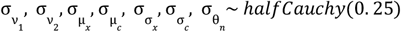

Posterior estimates were obtained using the NUTS-sampler, as implemented in pymc 5, with a target acceptance rate of 90%. We obtained 4 chains with 3000 samples each (1500 burnin). If we encountered divergences, we increased the target acceptance rate to 92.5%. To obtain estimates of the size and presence of an effect, we used “Bayesian” p-values –the probability mass of the posterior estimate above/below 0^126^. We generally used one-tailed cutoffs of 5% probability mass.

Model comparisons were performed using the estimated log pointwise predictive density (ELPD^48^) which is a state-of-the art model comparison technique that – unlike other information theoretic measures like BIC and DIC – takes into account the shape of the posterior distribution and the *effective* number of parameters of the model.

## MRI parameters

### 3T

We acquired functional MRI data using the Philips Achieva 3T whole-body MR scanner equipped with a 32-channel MR head coil, located at the Laboratory for Social and Neural Systems Research (SNS-Lab) of the UZH Zurich Center for Neuroeconomics. We collected 8 runs of fMRI data with a T2^*^-weighted gradient-recalled echo-planar imaging (GR-EPI) sequence (150 volumes + 5 dummies; flip angle 90 degrees; TR = 2286 ms, TE = 30ms; matrix size 96 × 70, FOV 240 × 175mm; in-plane resolution of 2.5 mm; 39 slices with thickness of 2.5 mm and a slice gap of 0.5mm; SENSE acceleration in phase-encoding direction (left-right) with factor 1.5; time-of-acquisition 4:52 minutes). Additionally, we acquired high-resolution T1weighted 3D MPRAGE image (FOV: 256 × 256 × 170 mm; resolution 1 mm isotropic; Shot TR = 2800ms; TI = 1098.6 ms; 256 shots, flip angle 8 degrees; TR = 8.3 ms; TE = 3.9 ms; SENSE acceleration in left-right direction 2; time-of-acquisition 5:35 minutes).

### 7T

We also acquired MRI data using a Philips Achieva 7T whole-body MR scanner (located at the Institute for Biomedical Technology of ETH Zurich) equipped with: 7T MRI quadrature transmit/32-channel head receive array coil (Nova Medical). For the functional data, we acquired T2^*^-weighed GRE-EPI sequences (150 volumes + 5 dummies); flip angle 74 degrees, TR=2300 ms; TE = 15ms; matrix size 109 x 128 (LR x PA); FOV 190 x 224 mm; in-plane resolution of 1.75 mm; 76 slices with thickness of 1.75 mm with no slice gap; SENSE-acceleration in phase-encoding direction (left-right) with factor 3; Multiband acceleration with factor 2; time-of-acquisition 5:35 minutes). At 7T, we acquired a T1-weighted 3D MPRAGE image with the following parameters: FOV = 240 × 240 × 160 mm; voxel resolution = 0.8 × 0.8 × 0.8 mm; inversion time (TI) = 1656.2 ms; 164 shots; flip angle = 7°; repetition time (TR) = 9.8 ms; echo time (TE) = 4.5 ms; SENSE acceleration factor = 3 (left-right) and 1.5 (anterior-posterior); total acquisition time = 4 minutes 43 seconds.

## fMRI preprocessing

Preprocessing on fMRI data was performed using fMRIPrep 20.2.2 ^127^, which is based on Nipype 1.6.1 ^128,129^.

### Anatomical data preprocessing

The T1-weighted (T1w) images collected from the 3T and 7T data were processed together. All of them were corrected for intensity non-uniformity (INU) with N4BiasFieldCorrection ^130^, distributed with ANTs 2.3.3^131^. The T1w-reference was then skull-stripped with a Nipype implementation of the antsBrainExtraction.sh workflow (from ANTs), using OASIS30ANTs as target template. Brain tissue segmentation of cerebrospinal fluid (CSF), white-matter (WM) and gray-matter (GM) was performed on the brain-extracted T1w using fast (FSL 5.0.9^FSL^ ^5.0.9;,132^). A T1w-reference map was computed after registration of 3 T1w images (after INU-correction) using mri_robust_template(FreeSurfer 6.0.1^133^). Brain surfaces were reconstructed using recon-all (FreeSurfer 6.0.1^134^), and the brain mask estimated previously was refined with a custom variation of the method to reconcile ANTs-derived and FreeSurfer-derived segmentations of the cortical gray-matter of Mindboggle^135^. Volume-based spatial normalization to one standard space (MNI152NLin2009cAsym) was performed through nonlinear registration with antsRegistration (ANTs 2.3.3), using brain-extracted versions of both T1w reference and the T1w template. The following template was selected for spatial normalization: ICBM 152 Nonlinear Asymmetrical template version 2009c^136^,

#### Functional data preprocessing

For each of the 24 BOLD runs found per participants (across all tasks and sessions), the following preprocessing was performed. First, a reference volume and its skull-stripped version were generated using a custom methodology of fMRIPrep. BOLD runs were slice-time corrected using 3dTshift from AFNI 20160207^137^. Head-motion parameters with respect to the BOLD reference (transformation matrices, and six corresponding rotation and translation parameters) are estimated before any spatiotemporal filtering using mcflirt (FSL 5.0.9^138^). A B0-nonuniformity map (or fieldmap) was estimated based on two (or more) echo-planar imaging (EPI) references with opposing phase-encoding directions, with 3dQwarp^137^ (AFNI 20160207). Based on the estimated susceptibility distortion, a corrected EPI (echo-planar imaging) reference was calculated for a more accurate co-registration with the anatomical reference. The BOLD reference was then co-registered to the T1w reference using bbregister (FreeSurfer) which implements boundary-based registration^139^. Co-registration was configured with six degrees of freedom. The BOLD time-series (including slice-timing correction when applied) were resampled onto their original, native space by applying a single, composite transform to correct for head-motion and susceptibility distortions. These resampled BOLD time-series will be referred to as preprocessed BOLD in original space, or just preprocessed BOLD. The BOLD time-series were resampled onto the following surfaces (FreeSurfer reconstruction nomenclature): fsaverage, fsnative. Several confounding time-series were calculated based on the preprocessed BOLD: framewise displacement (FD), DVARS and three region-wise global signals. FD was computed using two formulations following Power 2014(absolute sum of relative motions^140^) and Jenkinson (relative root mean square displacement between affines^138^). FD and DVARS are calculated for each functional run, both using their implementations in Nipype (following the definitions by Power et al. 2014). The three global signals are extracted within the CSF, the WM, and the whole-brain masks. Additionally, a set of physiological regressors were extracted to allow for component-based noise correction(CompCor^141^). Principal components are estimated after high-pass filtering the preprocessed BOLD time-series (using a discrete cosine filter with 128s cut-off) for the two CompCor variants: temporal (tCompCor) and anatomical (aCompCor). tCompCor components are then calculated from the top 2% variable voxels within the brain mask. For aCompCor, three probabilistic masks (CSF, WM and combined CSF+WM) are generated in anatomical space. The implementation differs from that of Behzadi et al. in that instead of eroding the masks by 2 pixels on BOLD space, a mask of pixels that likely contain a volume fraction of GM is substracted from the aCompCor masks. This mask is obtained by dilating a GM mask extracted from the FreeSurfer’s aseg segmentation, and it ensures components are not extracted from voxels containing a minimal fraction of GM. Finally, these masks are resampled into BOLD space and binarized by thresholding at 0.99 (as in the original implementation). Components are also calculated separately within the WM and CSF masks. For each CompCor decomposition, the k components with the largest singular values are retained, such that the retained components’ time series are sufficient to explain 50 percent of variance across the nuisance mask (CSF, WM, combined, or temporal). The remaining components are dropped from consideration. The head-motion estimates calculated in the correction step were also placed within the corresponding confounds file. The confound time series derived from head motion estimates and global signals were expanded with the inclusion of temporal derivatives and quadratic terms for each ^142^. Frames that exceeded a threshold of 0.5 mm FD or 1.5 standardised VARS were annotated as motion outliers. The BOLD time-series were resampled into standard space, generating a preprocessed BOLD run in MNI152NLin2009cAsym space. First, a reference volume and its skull-stripped version were generated using a custom methodology of fMRIPrep. All resamplings can be performed with a single interpolation step by composing all the pertinent transformations (i.e. head-motion transform matrices, susceptibility distortion correction when available, and co-registrations to anatomical and output spaces). Gridded (volumetric) resamplings were performed using antsApplyTransforms (ANTs), configured with Lanczos interpolation to minimize the smoothing effects of other kernels ^143^. Non-gridded (surface) resamplings were performed using mri_vol2surf (FreeSurfer). First, a reference volume and its skull-stripped version were generated using a custom methodology of fMRIPrep. BOLD runs were slice-time corrected using 3dTshift from AFNI 20160207^137^. The BOLD reference was then co-registered to the T1w reference using bbregister (FreeSurfer) which implements boundary-based registration^139^. Co-registration was configured with six degrees of freedom. The BOLD time-series (including slice-timing correction when applied) were resampled onto their original, native space by applying the transforms to correct for head-motion. These resampled BOLD time-series will be referred to as preprocessed BOLD in original space, or just preprocessed BOLD. The BOLD time-series were resampled onto the following surfaces (FreeSurfer reconstruction nomenclature): fsaverage, fsnative. The BOLD time-series were also resampled into standard space, generating a preprocessed BOLD run in MNI152NLin2009cAsym space. Many internal operations of fMRIPrep use Nilearn 0.6.2^144^, mostly within the functional processing workflow. For more details of the pipeline, see the section corresponding to workflows in fMRIPrep’s documentation.

## fMRI analyses

The main goal of our fMRI analyses was to get a trialwise estimate of the acuity of the representation of payoff magnitudes in the numerical parietal cortex^35^. Following earlier work^35,40,41^, we used an encoding/decoding-modelling approach, where we inverted an encoding model that describes how a voxel *i* responds to specific stimulus magnitude *s f*(*s*)→*x*_*i*_ in a Bayesian framework, by extending the encoding model to a multivariate likelihood function *p*(*X*|*s*) using a multivariate t-distribution: [*x*_1_, .., *x*_*n*_] ~ *f*_1..*n*_ (*s*) + ϵ with ϵ ~ *t*(0, Σ, *d*), where Σ is the residual covariance and *d* is the degrees-of-freedom of the t-distribution. Using an explicit likelihood function allows us to decode from trial-to-trial what was the presented payoff magnitude, as well as the acuity of the neural response. For every trial, we test the consistency of the BOLD activation pattern for all possible numerosities. The acuity of the neural response was operationalised as the dispersion (standard deviation) of the decoded posterior. Loosely speaking, the posterior will be less dispersed when the BOLD responses of many voxels agree with a small set of numerosities. When the amplitude of the response across voxels is smaller and/or they are not consistent with the same numerosities, the posterior will be more dispersed, as the decoder is less certain about which stimulus was presented. We postulate, as other have done before^40,41^, that the uncertainty of our decoding of the presented stimulus is related to the acuity of the *neural representation* of that stimulus.

The main fMRI analysis can roughly be split up in the following steps: 1) Fit a single-trial GLM to estimate trialwise measures of the response amplitude across voxels 2) Fit a numerical receptive field model^43^ to the response, 3) Fit a multivarite noise model to the residuals of the nPRF model in a leave-one-run-out cross-validation scheme^41^ 4) Obtain a posterior estimate of the payoffs magnitudes of unseen data using the noise model and an inverted nPRF model.

### Single trial estimates

We used the GLMSingle Python package^145^ to obtain single-trial BOLD estimates. Briefly, the GLMSingle package uses cross-validation to do model selection over GLMs with a) a library of different hemodynamic response functions b) different L2-regularisation parameters to shrink the single trial estimates, combating the issue of correlated single trial regressors^146^. c) GLMSingle also obtains GLMDenoise^147^ regressors based on the first *n* PCA components in a set of noise voxels. Noise voxels are defined as having low explained variance in the task-based GLM. The number of PCA components is selected via cross-validation.

As input to GLMSingle, we used both the 24 first and 24 second payoff presentations per run. For the second payoff pesentations, we modeled all trials with the same numerosity as being in the same condition, to aid GLMSingle with cross-validation (similar numerosities should have similar responses). After extensive analysis piloting, we chose to not include any additional confound regressors (e.g. motion parameters, RETROICOR parameters or aCompCorr regressros) in addition to the GLMDenoise confound regressors, as additional regressors did not lead to robustly increased (or even decreased) decoding accuracy and GLMDenoise. This is in line with earlier work on GLMDenoise^147^ and the related aCompCorr approach^141^.

### Numerical receptive field modelling

We fitted a numerical receptive field model to all voxels in the brain, for each session separately (so using 192 single trial estimates). The method is described in detail elsewhere^35,43^, so we only briefly describe it here. First, we estimate the mean µ, standard deviation σ, amplitude *A* and baseline *B* of a log-normal receptive field for every voxel in the brain separately, to predict the BOLD response of these voxels to different numerosities.

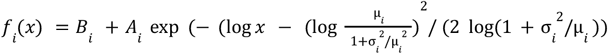

The second part of this equation is a parameterisation of the log-normal probability density function where µ and σ are the mean and standard deviation of the distribution in natural space. Although this parameterisation is somewhat exotic, it is highly useful when plotting the estimated preferred numerosities and their dispersion, for example on the cortical surface.

We first fit the model by using a grid-search: We correlatethe single trial estimates for the first payoff stimulus presentation with the predictions of a large grid of 60 µ-s between 5 and 80 and 60 σ-s between 5 and 40. We then estimate *I* and *A* linear least-squares on the best-correlating µ and σ-parameters. Finally, we used gradient descent^148^ to refine parameters further. The explained variance *R*^2^ for each voxel is then transformed to *fsaverage*-surface space to compare across participants. We compared the *R*^2^-maps to an ROI of the (right) numerical parietal cortex (rNPC) in *fsaverage* we drew for an earlier study. We observed that the *R*^2^-maps were tightly related to the (see Figure 2) and therefore chose to use this rNPC-mask in all follow-up analyses. The rNPC mask was transformed from *fsaverage*-surface space to individual anatomical space using *neuropythy* (https://github.com/noahbenson/neuropythy).

### Voxel selection

One key methodological choice in the encoding/decoding-framework is which voxels to include in the decoding step. In earlier work we chose an arbitrary number of voxels/surface vertices^35^. Although the number of voxels seemed to make little difference in individual decoding accuracy, it remains an arbitrary choice. Here, we chose to use a nested cross-validation scheme to select the number of voxels for each participants/run in a principled way. The method takes into account both that noisy voxels should have an out-of-sample *R*^2^ lower than 0 and different brains have different sizes. Specifically, when decoding run *i* (out of a total of 8 runs per session), we would fit the nPRF model on all voxels within the NPC mask on (6-run) subsets of the remaining 7 runs, by leaving another run *j* out *R*__*i*_*j*_. Then, for each of these nested folds *R*__*i*_*j*_, we calculated the *out-of-sample R* ^2^on run *j* for each voxel. Thus, for each voxel and each decoded test run *i*, we had 6 out-of-sample *R*^2^ *s*, which we averaged over. For decoding run *i*, we would only use voxels that had a mean out-of-sample *R*^2^ larger than 0.

### Decoding

After voxel selection, we used a leave-one-run-out cross validation scheme where the nPRF model was fitted to all runs but the test run, after which also a multivariate noise model was fitted to the residual signals. Specifically, we fitted the following covariance matrix:

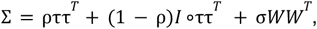

as well as the degrees-of-freedom of a 0-centred, multivariate t-distribution. Briefly, τ is a vector as its length the number of voxels (*n*) in the ROI. It pertains to the standard deviation of the residuals of each voxel. Thus, ρ determines to which extent all voxels correlate with each other ( ττ^*T*^ is the covariance matrix of perfectly correlated voxels, whereas *I* ∘ττ^*T*^ corresponds to a perfectly diagonal matrix/spherical covariance). *W* is a square matrix of *n* ×*n*. Each element *W*_*i,j*_ is the product of the receptive fields of voxels *i* and *j* across stimulus space (*f*_*i*_ (*S*) *f*_*j*_ (*S*) ^*T*^ with *S* being the entire stimulus space [5, 6, .., 112]. Thus, the free scalar parameter σ determines to which extent voxels with overlapping receptive fields have more correlated noise.

Once the noise model is fitted, we can now determine a likelihood function for any multivariate BOLD pattern *X* and for any numerical stimulus *s*:

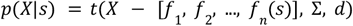

Since we assume a flat prior on all integers between 5 and 112, we can now just evaluate this likelihood on all these integers and normalize the resulting probability mass function (pdf) to integrate to 1. We take the expected value of this pdf to be our estimate of the presented stimulus. We take the standard deviation of the pdf as a measure of the acuity of neural coding. This standard deviation is used to split up trials within participants and payoff-numerosities for the probit model and as input to the PMCM as a linear regressor on all the model parameters.

## Acknowledgements

We are grateful to C. Schnyder, K. Treiber and M. Moisa at the Zurich Center for Neuroeconomics and Roger Lüchinger at the IBT for their excellent assistance in scanning, recruitment and participant facilitation. C.C.R. received funding from the University Research Priority Program ‘Adaptive Brain Circuits in Development and Learning’ at the University of Zurich and the Swiss National Science Foundation SNSF (grant no. 100019L-173248). G.d.H.was funded by the Dutch Research Council NWO (Rubicon grant no. 019.183SG.017/8O3B) and the University of Zurich (Forschungskredit grant no. K-33153-02-01)

## Code availability

All the code used in this package can be found online on github: https://github.com/Gilles86/risk_experiment/

Our computational cognitive models are implemented in the open-source Python library *bauer* https://github.com/ruffgroup/bauer/tree/main

The used nPRF model and decoding software algorithms are implemented in the open-source Python library *braincoder* https://braincoder-devs.github.io/

## Footnotes

1 https://github.com/ruffgroup/bauer/

