## Supplementary Materials for "Rapid Changes in Risk Attitudes Originate from Bayesian Inference on Parietal Magnitude Representations"

### Supplementary Figures

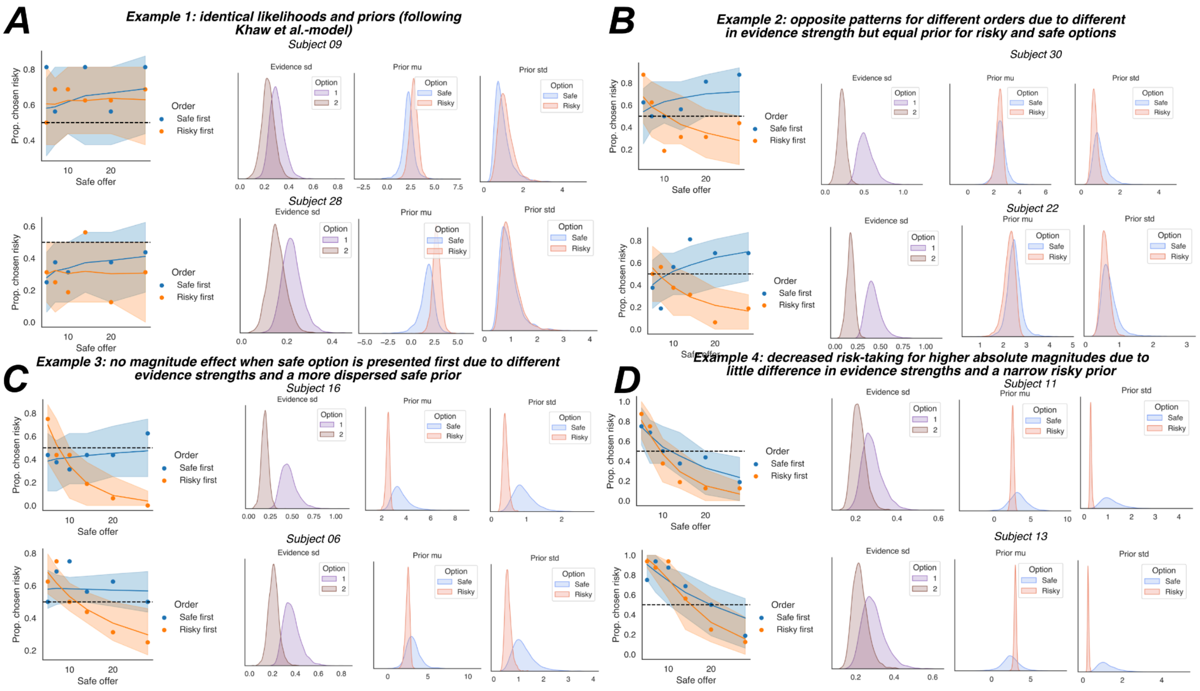


**Supplementary figure 1:** Four examples of qualitatively different behavioural patterns found in empirical data and how they are explained by our model parameters. **A)** Some participants show (practically) identical noise on the payoffs for the two options and use a common prior for both risky and safe payoffs. Therefore, they behave according to the KLW-model and are not influenced by the absolute magnitudes of the options, nor the order in which they were presented. **B)** Other participants have noisier representations for the payoff of the first option versus the second option, but appear to employ a common prior for the risky and safe option. Therefore, they show increasingly risk-seeking behaviour for larger absolute magnitudes when the safe option was presented first, but show the opposite pattern when the risky option came first. **C)** A third group of participants shows both a noisier representation of the first-presented option's payoff and different priors for the safe and risky option. The prior for the safe payoffs is generally more dispersed (larger sd), making the regression effects much more pronounced when the risky option was first presented versus when the safe option was presented first. **D)** A fourth group of participants showed little difference in noise for the first and second-presented options, but do show a much more dispersed prior for the safe option. These participants show more risk-averse behaviour for larger magnitudes, independent from the order in which the options were presented.

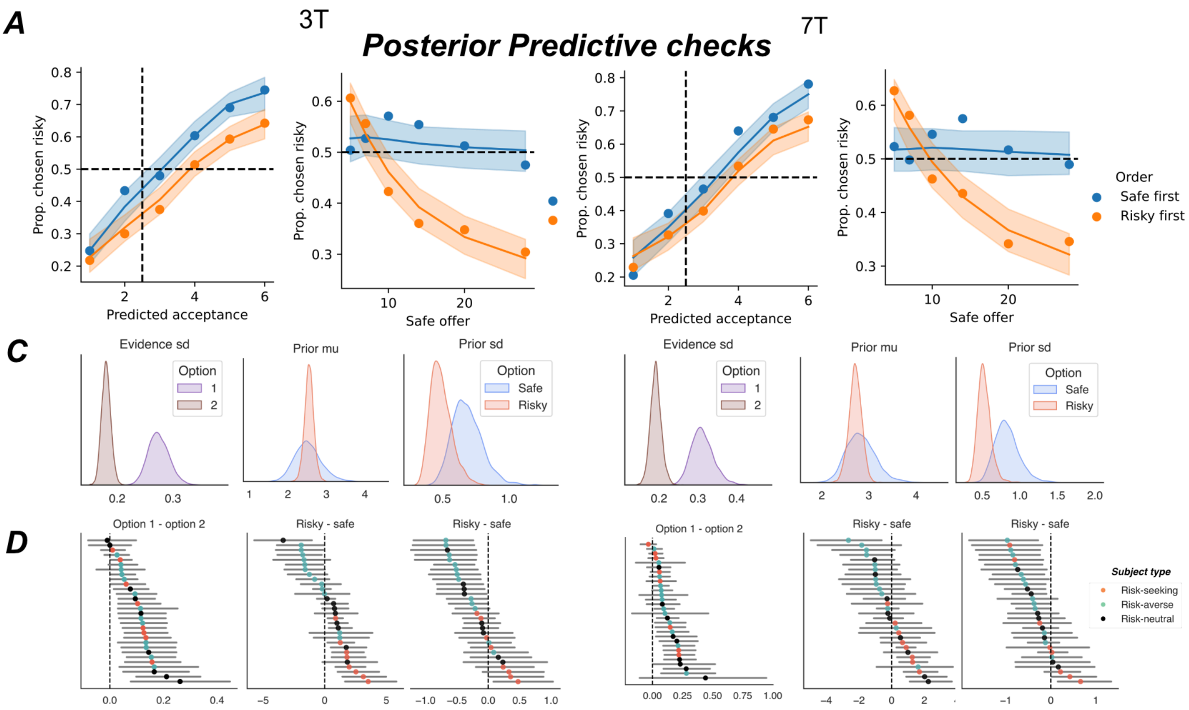


**Supplementary figure 2: Analysis of the KLW model and the extended version, separately for the two MRI sessions, analogous to Figure 2 in the main text.**

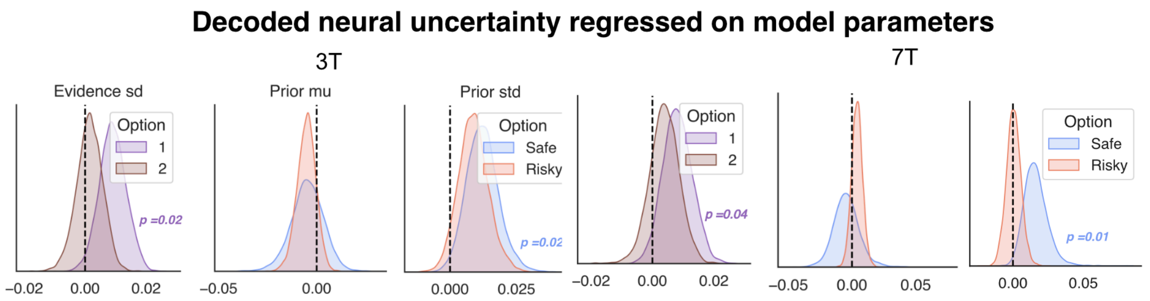


**Supplementary figure 3: analysis of the impact of decoded neural uncertainty on risk attitude, seperately for two sessions.** Posterior estimates of regression coefficients of trialwise decoded neural uncertainty on the 6 main parameters of the process model.

### Supplementary Texts

#### Supplementary text 1

#### Parameter Recovery study

In this study, we introduce a new model of risky choice with a relatively large number of parameters. Therefore, it is crucial to confirm that these parameters can be recovered from data[1,2]. As a first test, we estimated model parameters for the 3T and 7T sessions separately, hypothesizing that if the model is recoverable, these parameters should correlate across sessions. We indeed find that all six parameters correlate significantly across participants, between sessions (see Table S3.1)

| parameter | r | CI95% | p-val | BF10 |
|---|---|---|---|---|
| n1_evidence_sd | 0.7141877675 | [0.48 0.85] | 9.32E-06 | 2613.583 |
| n2_evidence_sd | 0.593567734 | [0.3 0.79] | 0.0005450728872 | 67.711 |
| risky_prior_mu | 0.4119552859 | [0.06 0.67] | 0.02369736843 | 2.609 |
| risky_prior_std | 0.7635675188 | [0.56 0.88] | 9.19E-07 | 2.15E+04 |
| safe_prior_mu | 0.5787956187 | [0.28 0.78] | 0.0008056913158 | 47.922 |
| safe_prior_std | 0.5527793204 | [0.24 0.76] | 0.001535733646 | 27.166 |

**Table S1.1: Correlation between mean parameter estimates of the 3T and 7T session, across participants.**

We also performed a parameter recovery study. We simulated data using the mean parameter estimates across the two sessions same number of trials and the same choice

problems as the participants in our study. Note that these choice problems were calibrated by a calibration session. We also performed a parameter recovery study where all 30 participants had identical choice problems (similar to García-Baretto et al.) and those results were nearly identical. We repeated the simulation-estimation cycle 10 times.
The raw results can be seen in Fig. S2.1.

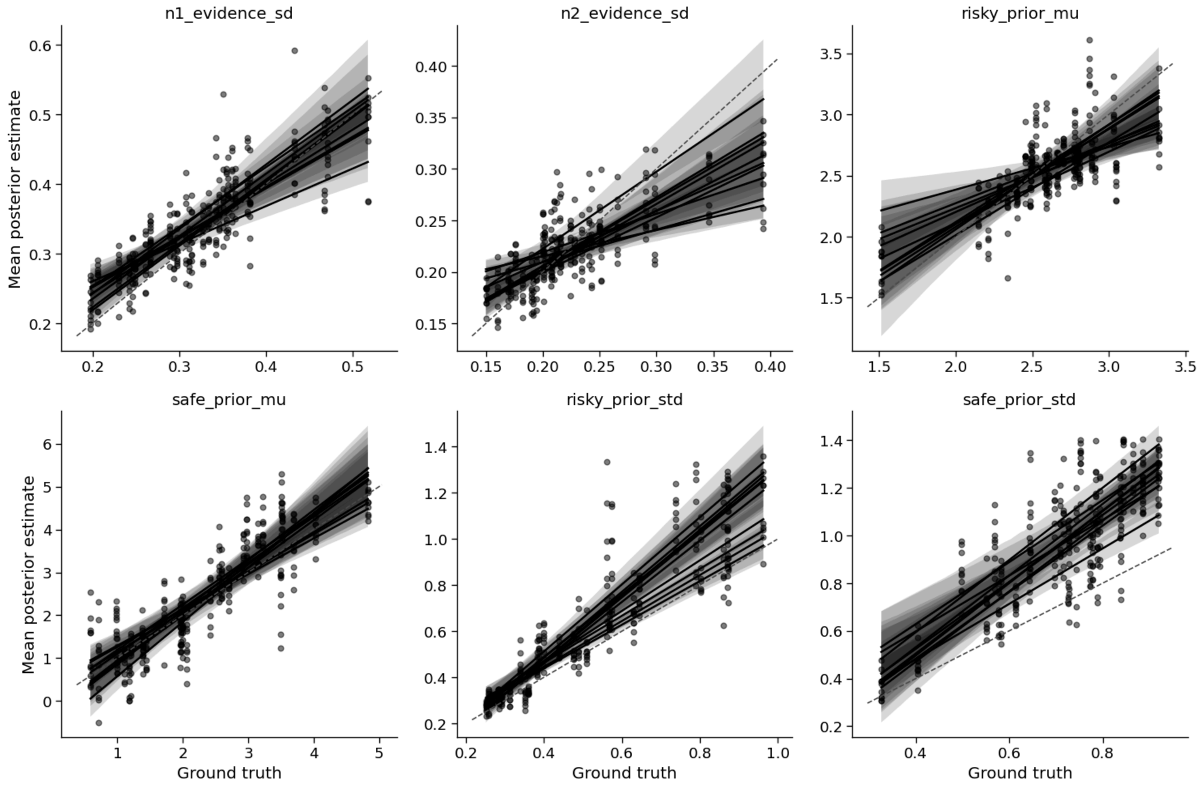


**Supplementary Figure S1.1:** Generating and estimated parameters for 10 simulations.

The figure suggests that all parameters are recovered. Figure S1.2. shows the average correlations between generating and estimated parameters.

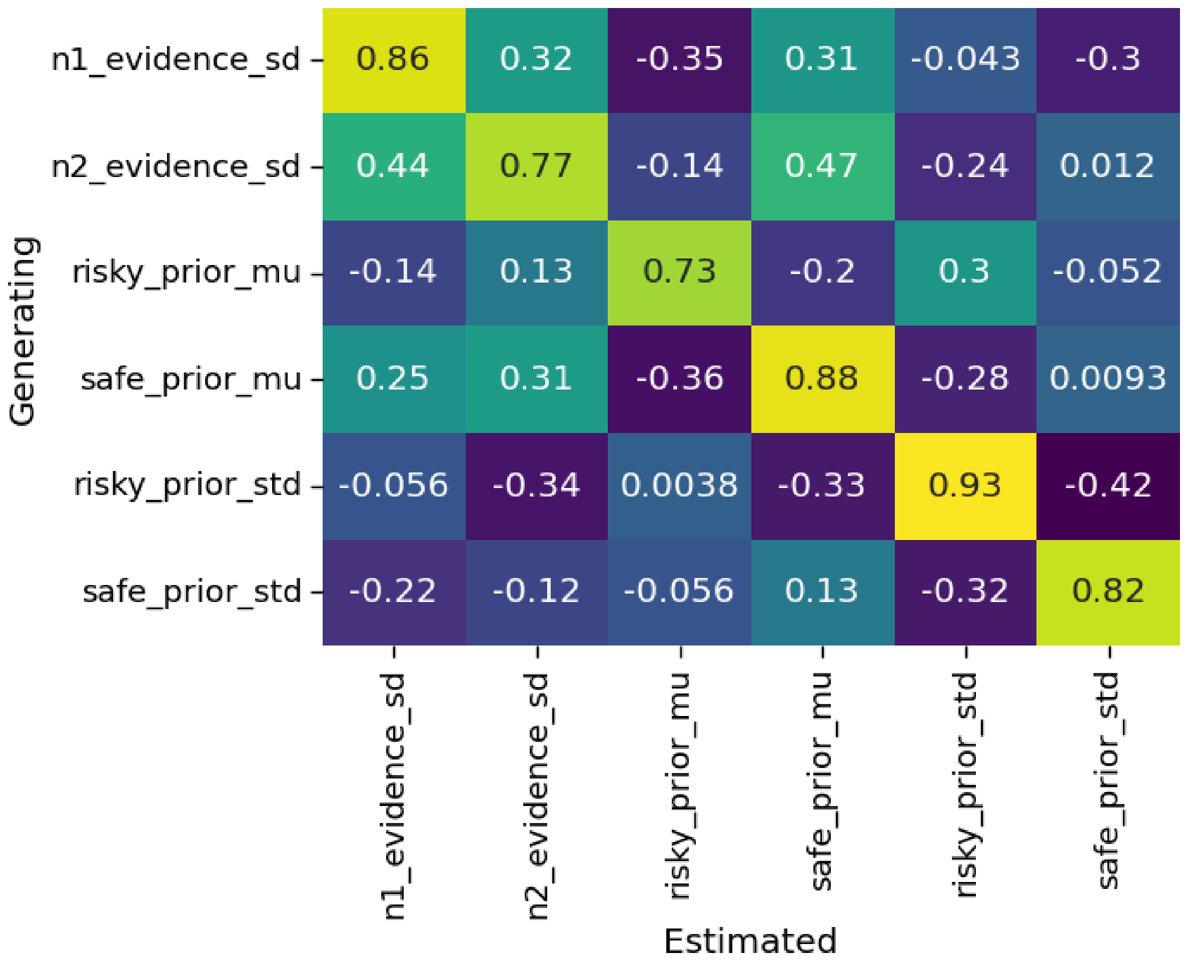


**Supplementary Figure S1.2:** Mean correlations between generating and estimated parameters.

The figure reveals that all estimated parameters are highly correlated with the *corresponding* generating parameters and much more so than with other parameters. However, some high correlations remain between estimated parameters and *different* generating parameters. E.g. the correlation between generating *n2_evidence_sd* and estimated *n1_evidence_sd* is 0.44. These correlations could be either due to a 'real' correlation in laten cognitive factors (which in this case seems quite plausible) but could also highlight a problem with parameter recoverability. Reassuringly, Supplementary Fig S1.3. shows that the off-diagonal correlations are generally already present in the generating parameters, and with similar magnitudes.

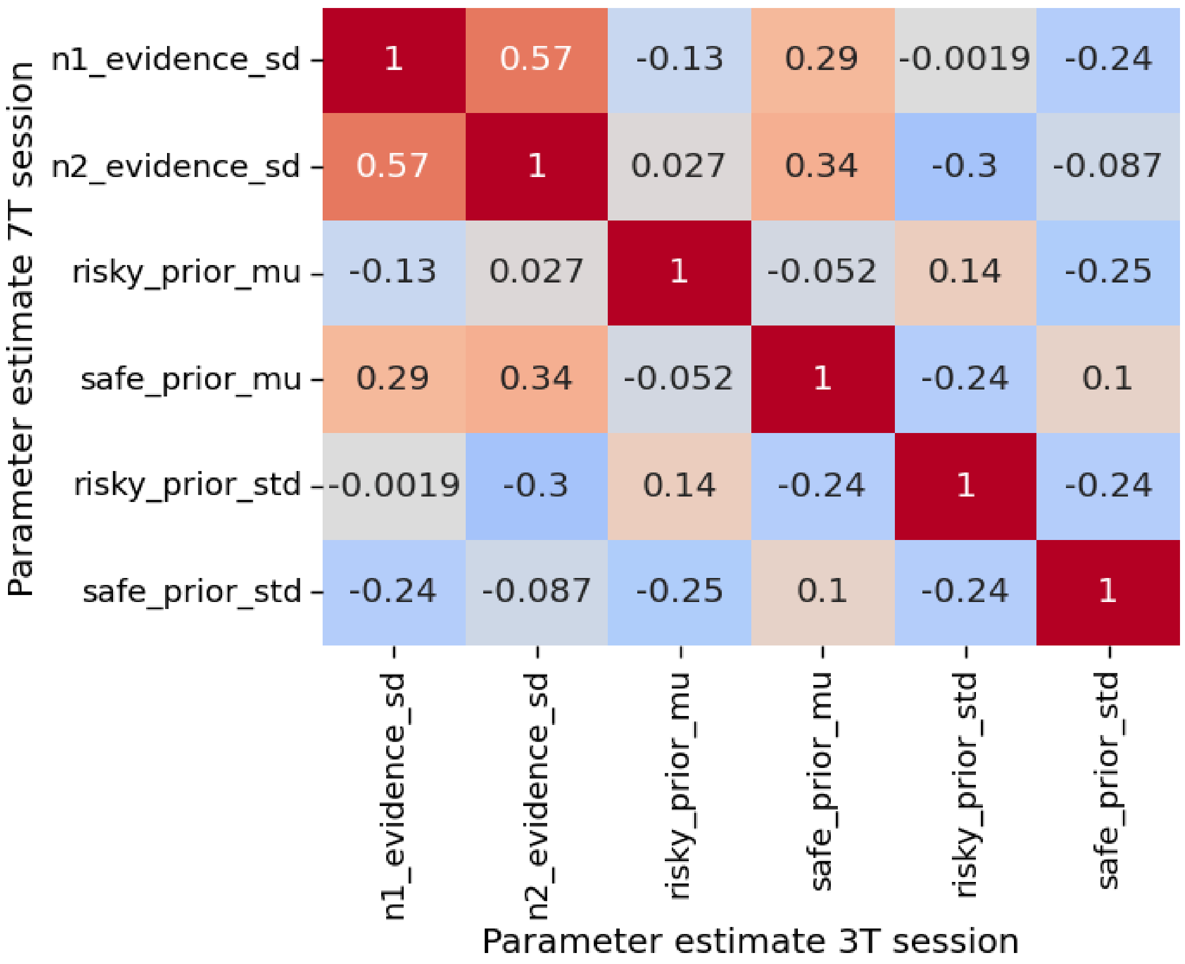


**Figure S1.3: Correlation matrix between (seperately estimated) mean posterior parameter estimates for session 3T an 7T**

To investigate this issue further, we also generated data from the prior distribution we used to estimate model parameter in real data (see methods section). The prior distributions assume independence (no correlation) between the six parameters in the population. We generated 100 samples from the prior distributions, simulated data for the same paradigm but without calibration and estimated them back using our hierarchical model We repeated this cycle 10 times. The mean resulting correlations can be seen in Figure S2.4.

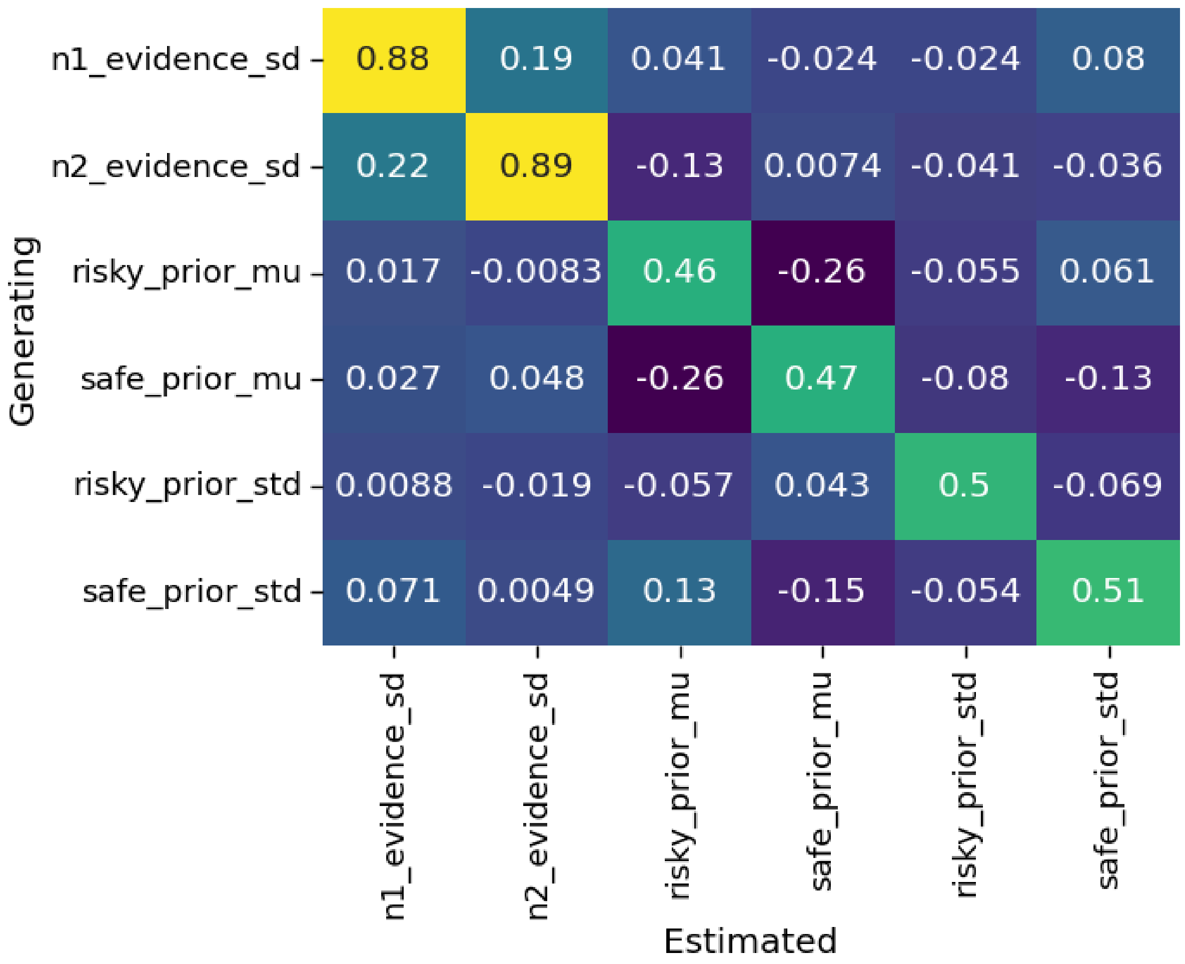


**Figure S1.4: Correlation matrix between generating and estimated parameters for parameter recovery using the prior samples as generating parameters**

Clearly, when the generating parameters are uncorrelated, the off-diagonal cells of the generating-estimation correlation matrix are much closer to 0. The correlations between estimated and generated parameters is also lower for the parameters that deal with prior beliefs. This is to be expected: When there is no difference in noise between the 1st and 2nd option, those parameters are not recoverable. Our prior distributions for n1_evidence_sd and n2_evidence_sd are, however, identical. Thus, for many samples, the difference between these two parameters is rather small (i.e., no working memory effect). In Figure S1.5, we plot generating and estimated parameters for different differences between *n1_evidence_sd* and *n2_evidence_sd*. Clearly, parameter recovery increases substantially when working memory are present. See also Table S1.2

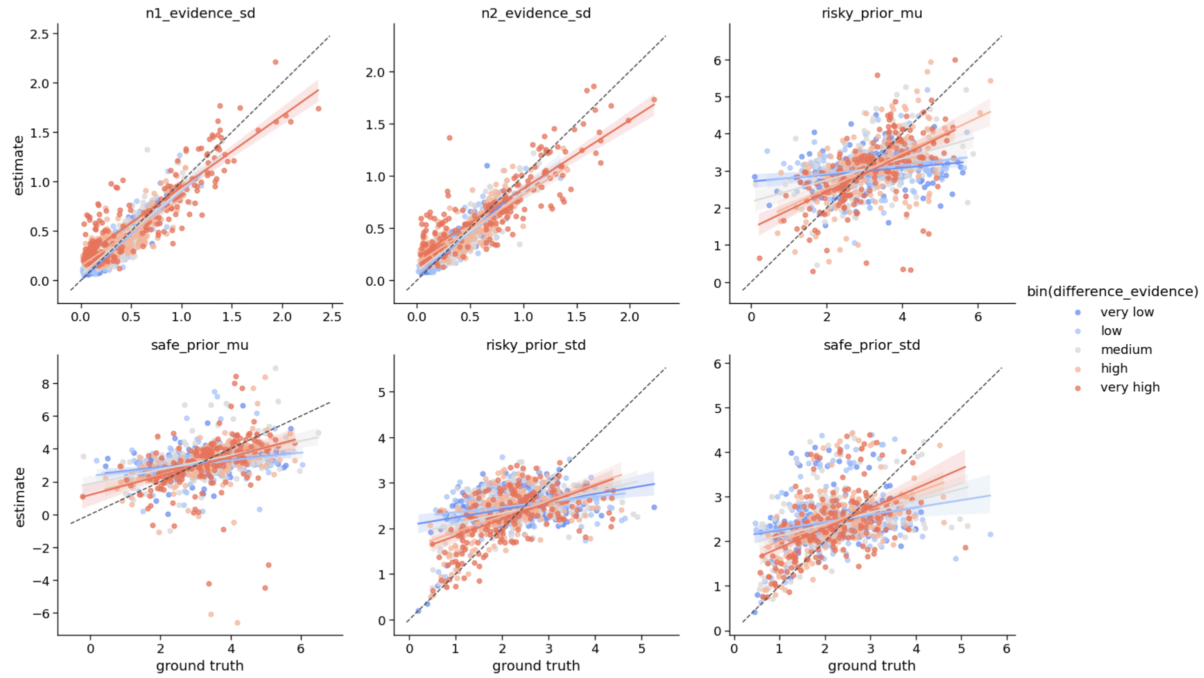


**Figure S1.5: Generated versus recovered parameters for the parameter recovery study for different sizes of the memory effect.**

| bin(difference_evidence) | n1_evidence_sd | n2_evidence_sd | risky_prior_mu | safe_prior_mu | risky_prior_std | safe_prior_std |
|---|---|---|---|---|---|---|
| very low | 0.932 | 0.901 | 0.214 | 0.315 | 0.361 | 0.256 |
| low | 0.873 | 0.879 | 0.229 | 0.268 | 0.272 | 0.214 |
| medium | 0.865 | 0.837 | 0.43 | 0.441 | 0.448 | 0.385 |
| high | 0.811 | 0.822 | 0.565 | 0.403 | 0.505 | 0.47 |
| very high | 0.878 | 0.863 | 0.563 | 0.371 | 0.553 | 0.508 |

**Table S1.2: Correlation between generating and mean estimated posteriors for different memory effect sizes**

In sum, our parameter recovery study shows that in general, all 6 parameters of the PMCM are identifiable. However, to which extent they can be recovered depends on the particulars of the parameter space that is sampled by the experiment. To be able to recover the different priors of risky and safe options, it is important that the noise in the two options orthogonally differs as well, e.g., through a memory effect as we introduced in our paradigm.

### Supplementary text 2

#### Model Recovery study

To ensure that we could distinguish between the 5 computational models we tested, we embarked on a model recovery study. Specifically, we (1) fitted all models to our empirical data (30 participants, 2 sessions), (2) simulated 100 data sets based on those parameters, (3) fitted our 5 models to all those simulated datasets, (4) used EPLD as a model comparison technique to select the best-fitting model.

As can be seen from Supplementary Fig. S2.1, the PMCM was correctly identified 100/100 times. Models where either the prior and noise are static were sometimes misrecovered, although the right model was picked in the large majority of cases.

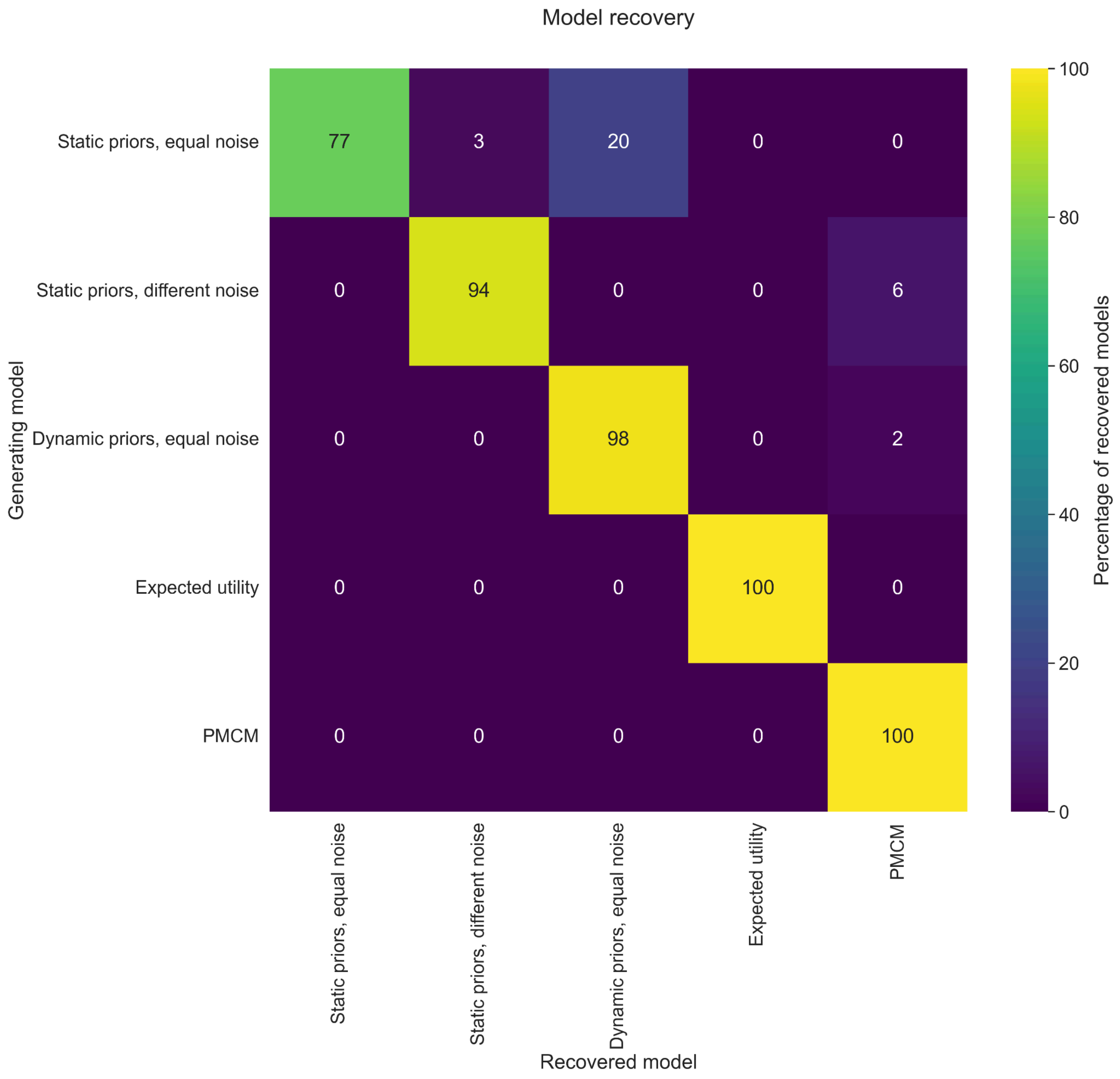

**Figure S2.1. Relative frequencies that models were recovered, conditional on generating models.**

### Supplementary Text 3: Order effect using symbolic payoffs (i.e., Arabic numerals)

#### Introduction

We ran an additional experiment to test whether the *order x stake*-effect we observed in the fMRI experiment –where participants were more likely to choose risky options when they were presented first and this effect increased for increased stake sizes– would generalize to a more common 'symbolic' presentation format using Arabic numerals for the payoff magnitudes. The main hypothesis was thus that also when using Arabic numerals as presentation format, earlier presentation of risky options would lead to relatively less risk taking, in particular when these payoffs were relatively large. We also hypothesized that this happened because the payoff that was presented first had to be kept in working memory and its representational acuity would therefore deteriorate.

#### Methods

Fifty-eight (n=58) participants (average age 24.6, range 20-41; 31 females) were recruited for the follow-up study. We informed them about the study's objectives, the data recorded and obtained from them, the tasks involved, and their expected payoffs. Our experiments conformed to the Declaration of Helsinki, and our protocol was approved by the Canton of Zurich's Ethics Committee. The experimental paradigm was highly similar to that of the fMRI experiment (see Figure S3.1A). The main difference with the fMRI experiment was that payoffs were not presented as dot clouds but using Arabic numerals and the experiment was performed in the behavioral laboratory, outside of the MRI scanner. Again, participants were instructed to choose between two prospects: one *safe* option, with a 100% probability of payout, and one *risky* option, with a 55% probability of payout. The safe options were uniformly sampled between 5 and 28 CHF *in log space*, and then rounded to the nearest cent in natural space (mean CHF 13.38, std. 6.49). The risky options were sampled by multiplying the safe option by a uniformly sampled number between 1 and 4 (mean CHF 33.35, std. 20.61). Crucially, in half of the trials, safe options were presented first, and in the other half risky options were presented first. Each subject performed 256 trials, presented across 8 runs of 32 trials. Each run was divided in two blocks, where either the risky or the safe option was presented first. Between the first and second option, there was an inter-stimulus interval of 4-6 seconds. The order of blocks within runs was counterbalanced. The total experiment took approximately 1 hour.

### Results

#### Raw behavioral results

We predicted that risky options would be more likely to be chosen when they are presented second, in particular when they are smaller. Indeed, participants were significantly more likely to choose risky options when they were presented second (41.1%, sd. 40.6%) versus first (37.9%, sd. 39.5%), $F(1,57) = 17.16$, $p=0.00012$). However, this effect did not interact with the stake sizes (5 equal bins, $F(4, 228) = 0.019$, $p=0.99$). There was also no main effect of stake size ($F(4, 228) = 0.32$, $p=0.86$; also see Fig. S3.1B-C).

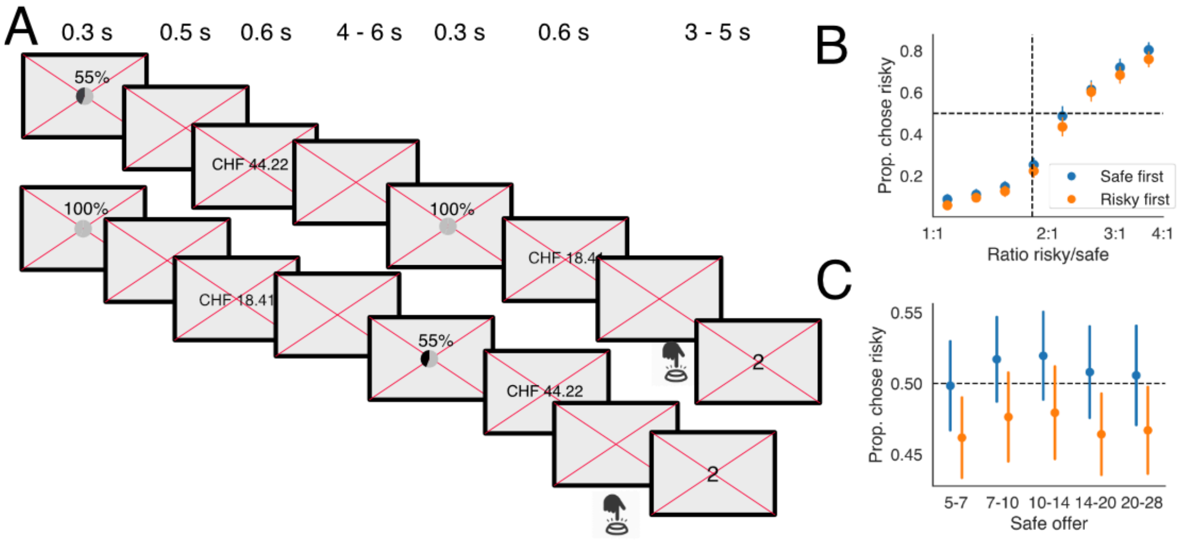


***Figure S3.1: A)*** *Participants were incentivized to choose between two prospects: A risky option with 55% probability of receiving the payoff and a safe option with 100% probability of receiving the payoff. Prospects were presented sequentially (4 to 6 seconds in between). Payoffs were presented as two-decimal amounts in CHF. Either the risky or the safe option was presented first and thus had to be held in working memory (all choice sets occurred twice for every order for every session).* ***B)*** *Average choice behaviour confirms that participants were more likely to choose the risky option when the safe option was presented first, in line with that assumption that options presented first (last) are noisier (less noisy) and thus more prone to a central tendency effect. Error bars correspond to the standard error of the mean over participants.* ***C)*** Average choice behavior for different stake sizes (defined as the quantile of safe offers). *Error bars correspond to the standard error of the mean over participants.*

#### Psychophysical modeling

When only analyzing average choice proportions, the effects of stake size and their interaction with order may be clouded by other factors, such as changes in choice consistency that come with different stake sizes. This is particularly likely for the presentation format using Arab numerals, where Weber's law clearly does not hold[3,4] and the choice sets for different stake sizes no longer have an equal psychophysical distance to the participant's indifference point. Therefore, we estimated a hierarchical, Bayesian psychophysical model[5,6]. We estimated four probit models, where the probability of choosing the risky option was always a function of the log-ratio of the risky and safe payoff, and, optionally, also 1) the magnitude of the safe option (5 bins) as a measure of overall stake size and 2) the order in which the options were presented (risky first or safe first). Formal model comparison using estimated pointwise predictive density (ELPD), which controls for model complexity[7], showed that a model where both order *and* overall magnitude were included outperformed all other models (See suppl. Table 1.1). This suggests that there is indeed an effect of both order, and magnitude, as well as an interaction between the two.

| Model | ELPD | est. n parameters | Model weight | dSE |
|---|---|---|---|---|
| order x stake | -5383.73 | 376.83 | 0.85 | 0 |
| stake | -5430.28 | 282.67 | 0.09 | 11.09 |
| order | -5595.52 | 161.92 | 0.07 | 22.01 |
| null | -5631.84 | 124.84 | 0 | 23.51 |

***Table S3.1: Model comparison psychophysical models including/excluding stake size and order**. ELPD: expected log pointwise predictive density. Model weight: weight according to stacking method. dSE: difference between top model and other models in standard errors of the ELPD.*

When we inspect the estimated indifference points of the winning model (Fig. S3.2), we can see that the difference in indifference points indeed increases with stake size. For the largest stake sizes (but not the smallest one), the 95% credible interval does not overlap with 0 anymore.

Finally, we replicate the finding in the main fMRI experiment, as well as earlier work[5] that the slope of the psychophysical function is positively correlated with the indifference point (defined as the risk-neutral probability) with $r(57)=0.34, p=0.008$. This again confirms that there is a close relationship between the noisiness of decision-makers and their deviation from risk-neutrality.

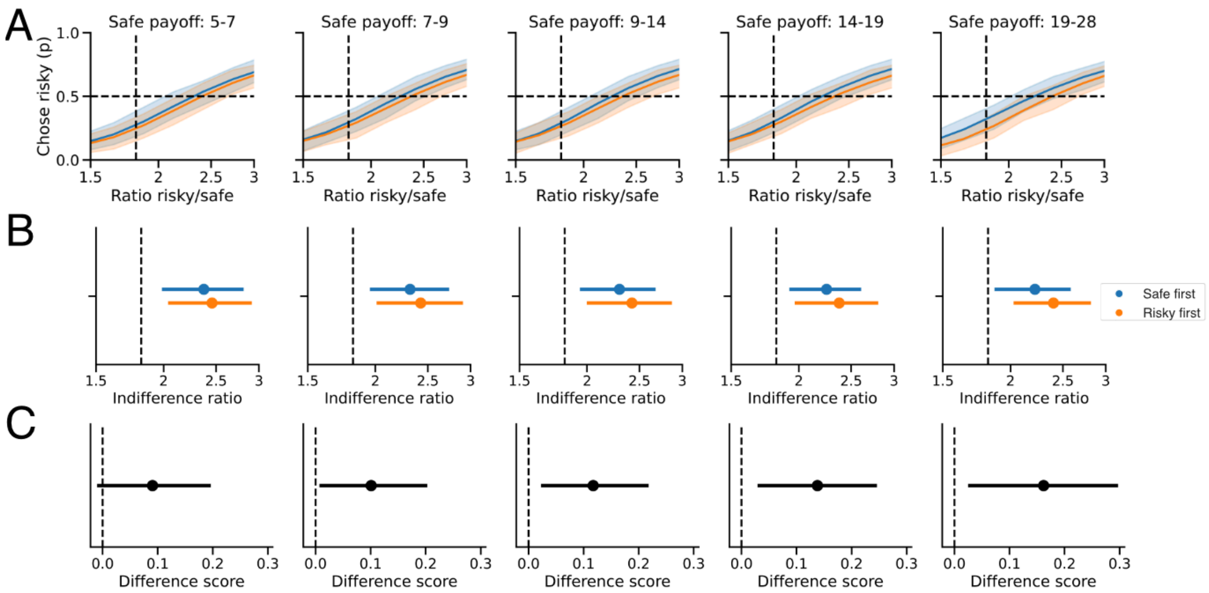


***Figure S3.2: A)** Mean estimated psychophysical curves: probability of choosing the risky option as a function of the log-ratio of the risk and the safe option. Different panels correspond to different stake sizes, different colors different orders of presentation.* ***B)*** *Estimated indifference points for different stake sizes and presentation orders. Note how the indifference points for trials where the safe option was presented have a lower ratio, which corresponds to less risk aversion.* ***C)*** *The difference of the indifference ratios for different stake sizes. Note how the difference increases as a function of stake size.*

In sum, the psychophysical modeling confirms the hypothesis that participants become more risk-averse when risky options are presented first. Moreover, this effect increases with stake size. This is consistent with a model where the fidelity of the representation of the payoffs of (risky) options that are kept in working memory increasingly deteriorates, showing larger distortions for larger payoff magnitudes.

#### PMC model

To get a more mechanistic understanding of the observed choice behavior and test whether it is consistent with increased noise for memorized options, we also fitted four different versions of our Perception and Memory-based Choice Model (PMC) model to the data. These four versions of the model differed on two dimensions: 1) whether the model assumed that participants could use different priors for the payoffs of the risky and safe options and 2) whether the model assumed different amounts of noise for the perception/representation of the first and the second option. We then used formal model comparison to test which model best explained the data, correcting for the increased number of parameters, using the ELPD[7].

Of the four tested models, the full PMC model (the one that also performed best for the fMRI data), where participants applied different priors and had differing amounts of noise for the first and the second option, performed best (see Table S3.2). Thus, these data suggest that, indeed, participants have differing amounts of noise for the first and second payoff, and they seem to integrate different prior beliefs about the potential payoffs of risky and safe options. Note that a standard expected utility model also gets outperformed by all other models.

| **Model** | **ELPD** | **est. n parameters** | **Model weight** | **dSE** |
|---|---|---|---|---|
| Different priors, different noise | -5376.19 | 249.87 | 0.79 | 0 |
| Different priors, equal noise | -5425.97 | 174.85 | 0.19 | 13.37 |
| Equal priors, different noise | -5669.45 | 192.04 | 0 | 23.94 |
| Equal priors, equal noise | -5701.00 | 118.02 | 0.02 | 26.45 |
| Expected utility model | -5795.26 | 156.44 | 0.1 | 36.71 |

***Table S3.2: Model comparison PMC models and expected utility model**. ELPD: expected log pointwise predictive density. Model weight: weight according to stacking method. dSE: difference between top model and other models in standard errors of the ELPD.*

In addition to the formal model comparison, we also performed a qualitative model comparison using posterior predictive plots (see Fig. S3.3). Note how only models that allow for different amounts of noise for the representation of the payoffs of the first and second option can convincingly capture the full order effect. The inclusion of different priors for risky and safe payoffs seems to be less important. The expected utility model fails to account for any order effect or interaction.

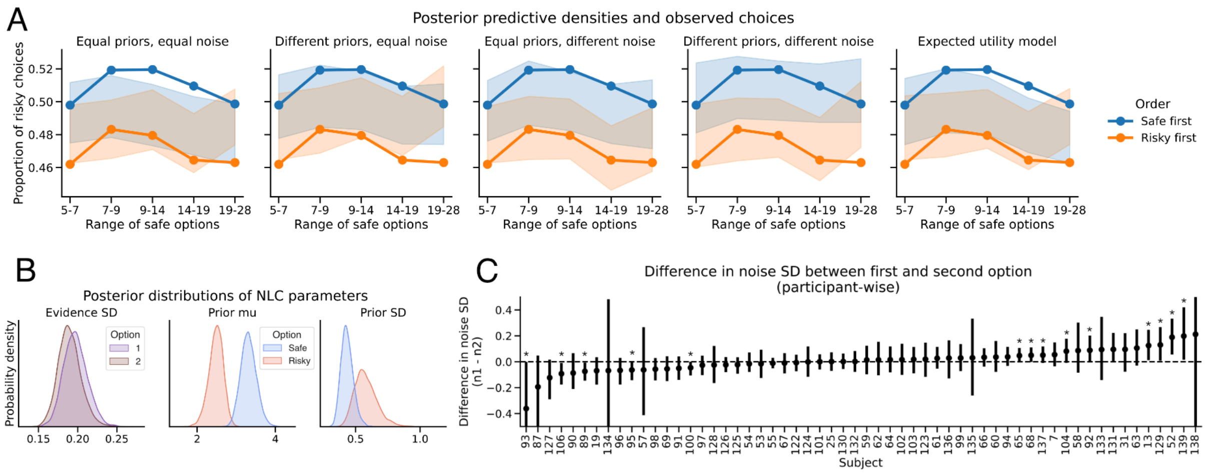


***Figure S3.3: A)*** *Posterior predictive checks for the different versions of the PMC model (four leftmost panels), as well as a standard expected utility model (rightmost panel). Note how only the PMC models that include varying noise between the first and second option can properly capture the order effect on choice proportions.* ***B)*** *Group parameter posteriors for the full PMC model: Evidence SD (or noisiness) for the first and second option, prior means for risky and safe options, as well as the dispersion of those two priors. Note how at the group level there is no robust difference in the average amount of noise between the first and second option.* ***C)*** *Individual estimates of the difference in noisiness between the first and second option. The leftmost subjects show a more noisy second option, whereas subjects on the right of the distribution show increased noise for the second option.*

The parameter estimates at the group level of the full PMC model (Fig. S3.3B) suggest that *at the group level*, there is no significant difference between the mean noisiness of the first and second payoff representation ($p_{Bayesian}$=0.37) –although the first option is estimated to be slightly noisier, as expected. Furthermore, on average, participants believe that risky options have relatively lower payoffs than safe options ($p_{Bayesian}$=0.0018) and the prior for risky options is wider than the one for safe options ($p_{Bayesian}$=0.0073).

At first glance, the finding of (approximately) equal noise for the 1st and 2nd option at the group level might seem at odds with the psychophysical results as well as the model comparison which suggest that only a model that allows for different noisiness of the first and second option representation can explain the patterns in the data. However, a closer inspection of the *individual* parameter estimates (Fig. S3.3C) suggests that there are large interindividual differences in the noisiness of the first versus second payoff representation. Out of 58 participants, 10 participants show a robustly more noisy representation of the first option (i.e., their 95% credible interval lies above 0), whereas 5 participants show a robustly more noisy representation of the second option compared to the first option. Roughly half of the participants show a difference very close to 0 (but note that we cannot use the credible interval to conclude that the difference is *equal* to 0).

Figure S3.4 illustrates the raw behavioral patterns of the 3 subgroups. Participants with noisier perception of the first option show the opposite pattern in terms of the interaction between stake sizes and order. Moreover, participants for which no robust difference in the noisiness with which the 1st and 2nd option get perceived can be found, at the group level still show a clear main effect of order (F(1, 42) = 9.26, p=0.004), suggesting these participants might still have a some working memory noise, but we lack statistical power to statistically establish this in each individual participant.

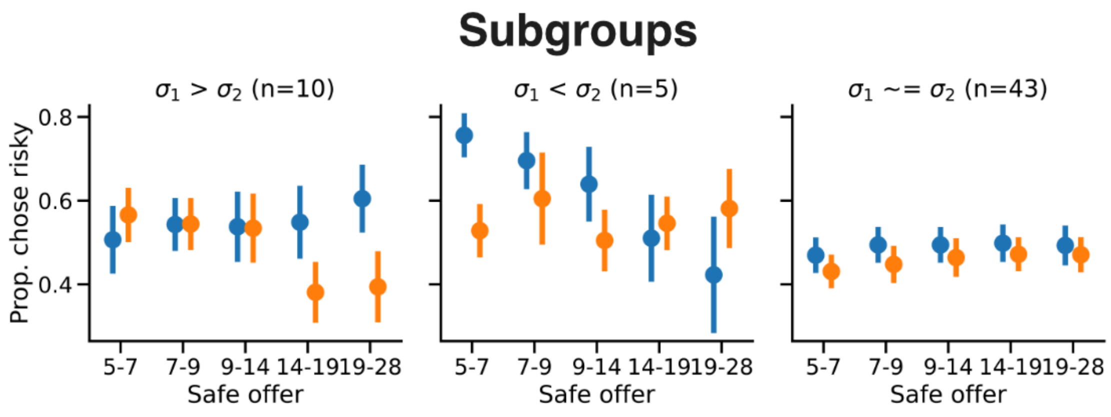


***Figure S3.4:*** *Different subgroups of participants for the symbolic task. About 10 participants showed a much noisier perception of the first payoff versus the 2nd payoff (left panel). For this group, both the Order (F(1, 9) = 10.04, p=0.011389 as the interaction between order (F(4, 36) = 8.94, p=0.000041) and stake size had a significant effect on choice proportions. Five (5) participants, however, showed a noisier perception of the second option compared to the first option, as if they decide immediately, based on the first option. These participants show no order effect (F(1,4), = 1.07, =0.36), but a very significant interaction effect (F(4, 16) = 11.05, p=0.00017). Finally, 43 participants showed roughly equal noise for the first and second option. However, these 43 participants did still show a significant order effect (F(1, 42) = 9.26, p=0.004), but no interaction effect (F(4, 168) = 0.22, p=0.92). Note that the statistical tests are for illustrative purposes only (revealing stereotypical patterns for certain parameter regimes) since the subgroups were selected based on the behavior that's tested. Error bars indicate standard error of the mean over subjects.*

#### Discussion

In this additional experiment, we set out to test whether the order in which risky prospects are presented impacts behavioral risk attitudes even when payoff magnitudes are presented in an unambiguous way, without any stimulus noise –namely as Arab numerals. We also speculated that, just as when payoffs are presented as stimulus clouds, participants would be particularly more risk averse when risky options presented first and when the average stakes were particularly large, because of increased downward central tendency effects. We found a very significant effect of order on the proportion of risky choices: Participants made more risk-averse choices when the risky options were presented first. Moreover, formal and qualitative model comparison revealed that this order effect increased as the stake sizes increased. This is consistent with an account where working memory noise impairs the neurocognitive representation of the first-presented option more than the second option.

We also fitted four different versions of our PMC model. Formal model comparison as well as inspection of posterior predictive densities suggest that only a full model, where participants have different beliefs about potential risky and safe payoffs, as well as different amounts of noise for the first and second option could explain the patterns in the empirical data. Parameter estimates, however, showed some surprising values. First, at the group level, there was no significant difference in the average amounts of noise for the representation of the first and second option. Conversely, at the level of individual parameters, a substantial group of participants (n=15) did show robustly more noise for the first *or* the second option (note that this is likely the case for more participants as well, but the parameter estimates at the individual level are relatively noisy). This suggests that order does play an important role for at least some participants, but they likely employ slightly different strategies. The obvious difference in strategy might be that some participants focus more on the first rather than the second option before they make their decision[8]. This result closely relates to work on the link between gaze and (economic) choice[9] and future work should further disentangle the effects of working memory and covert/overt attention[10].

The fact that the prior for risky options was estimated to have a lower mean than the prior for safe options is somewhat more puzzling. Our model (and earlier versions of it[5]) assume a linear utility function, they do not model actual risk preferences (over and above the effects of biased perception) in any way. We speculate that if participants are particularly risk-averse, the model might 'capture' this by subjective priors that do not exactly capture the objective priors. Another important issue that might explain the  is that the assumption of the PMCM (just as earlier models[5,11,12]) of constant noise in log space probably does not hold for Arab numerals (see probit results and, e.g., [4]), in particular not when the range of payoffs is more than one order of magnitude as was the case in this experiment. Future modeling effort should try to account for this issue, potentially by relaxing the noise scale invariance assumption (e.g., [4]).
